# Quantifying the propagation of parametric uncertainty on flux balance analysis

**DOI:** 10.1101/2021.09.16.460685

**Authors:** Hoang V. Dinh, Debolina Sarkar, Costas D. Maranas

**Affiliations:** Department of Chemical Engineering, Pennsylvania State University, University Park, Pennsylvania, USA; Center for Advanced Bioenergy and Bioproducts Innovation, The Pennsylvania State University, University Park, PA 16802, USA

## Abstract

Flux balance analysis (FBA) and associated techniques operating on stoichiometric genome-scale metabolic models play a central role in quantifying metabolic flows and constraining feasible phenotypes. At the heart of these methods lie two important assumptions: (i) the biomass precursors and energy requirements neither change in response to growth conditions nor environmental/genetic perturbations, and (ii) metabolite production and consumption rates are equal at all times (i.e., steady-state). Despite the stringency of these two assumptions, FBA has been shown to be surprisingly robust at predicting cellular phenotypes. In this paper, we formally assess the impact of these two assumptions on FBA results by quantifying how uncertainty in biomass reaction coefficients, and departures from steady-state due to temporal fluctuations could propagate to FBA results. In the first case, conditional sampling of parameter space is required to re-weigh the biomass reaction so as the molecular weight remains equal to 1 g/mmol, and in the second case, metabolite (and elemental) pool conservation must be imposed under temporally varying conditions. Results confirm the importance of enforcing the aforementioned constraints and explain the robustness of FBA biomass yield predictions.

## 1. Introduction

Flux balance analysis (FBA) encompasses a wide range of computational analyses that operate on genome-scale metabolic (GSM) models (Lewis et al., 2012; Orth et al., 2010), and provide insights on maximum theoretical yields of biomass or product per mole of substrate or limiting resource (Feist and Palsson, 2010), and metabolic activities in the form of intracellular fluxes under physiological constraints (Long et al., 2017; Mahadevan and Schilling, 2003). Relying on the solution of a linear programming (LP) optimization formulations (Orth et al., 2010), FBA analyses have gained broad acceptance due to the tractability and scalability of the calculations and the wide availability of GSM models for many organisms (Gu et al., 2019). As many as 185 manually curated GSM models have been reported (as of February 2019 (Gu et al., 2019)) with many thousands (Gu et al., 2019; Lieven et al., 2020) automatically generated using workflows such as KBase (Arkin et al., 2018), RAVEN (Wang et al., 2018), and CarveMe (Machado et al., 2018). GSM models aim to encompass all known biochemical conversions occurring in an organism, the participating metabolites, reaction stoichiometry, corresponding enzymes, genes encoding them, and production of biomass constituents in physiologically relevant proportions in cells (Thiele and Palsson, 2010). FBA predictions have been found to recapitulate growth phenotypes reasonably well under nutrient-limited conditions (O’Brien et al., 2013) and after adaptive laboratory evolution (Ibarra et al., 2002). By exploiting the fact that the timescale of organism growth far exceeds that of cellular biochemical reactions, every metabolite can be assumed to be at a pseudo-steady state, whereby the production equals the consumption flux. FBA also requires the establishment of an objective function as a teleological driver of cellular metabolism, which is usually maximizing biomass yield for a given amount of carbon uptake (Feist and Palsson, 2010). This translates computationally to maximizing flux through an imposed biomass drain (i.e., biomass reaction) which serves to sequester metabolites in their physiological ratios. Thus, at the heart of most FBA studies lie two major assumptions: (i) the biomass precursors and energy requirements are constant and invariant to changes in growth conditions, environmental, and/or genetic perturbations; and (ii) metabolites remain at steady-state with no temporal fluctuations in their concentration or production/consumption rates.

Uncertainty or inaccuracies in biomass composition can arise from limitations in measuring techniques (Beck et al., 2018; Lange and Heijnen, 2001), and the metabolically and temporally varying, and/or heterogenous nature of cellular populations (Takhaveev and Heinemann, 2018). Experimental quantification and modeling of metabolic heterogeneity is important for accurate cellular characterization (Takhaveev and Heinemann, 2018) and efficient bioprocessing (González-Cabaleiro et al., 2017). A previous metabolic modeling effort accounting for biomass composition variability considered biomass coefficients to be binomially distributed (MacGillivray et al., 2017). However, this effort did not enforce the requirement that the molecular weight (MW) of the biomass metabolite must always equal 1 gram dry weight (gDW) mmol^−1^ (Thiele and Palsson, 2010), which is needed to correctly map biomass flux to specific growth rate (Chan et al., 2017). In addition to the biomass composition, variance in energy demand parameters can be significant factors controlling predicted maximum theoretical yields. Growth-associated ATP maintenance (GAM) captures the energy demand per unit of produced biomass (Thiele and Palsson, 2010) and is embedded within the biomass reaction. Non-growth associated ATP maintenance (NGAM) accounts for the energy demand associated with cellular processes such as repair and maintenance (Thiele and Palsson, 2010). GAM and NGAM values are estimated using regression on growth data from chemostats (Thiele and Palsson, 2010), and are thus subject to uncertainty due to instrument limitations and population heterogeneity to an even larger extend than biomass composition. Capturing uncertainty in ATP maintenance parameters is essential because fluctuations in energy demand can propagate to other parts of the metabolic network and ultimately affect the growth and product yield.

The steady-state condition for all metabolites is a fundamental assumption of FBA. However, this might be too stringent as time-course intracellular metabolomics studies have revealed the transient nature of metabolite pools (Andreas Angermayr et al., 2016; Kultschar et al., 2019; Qiao et al., 2020; Yurkovich et al., 2017) as a cellular response to microenvironmental variations (Schmitz et al., 2017) and over the course of a typical growth cycle (Andreas Angermayr et al., 2016). In addition, periodic variations in metabolism are inherent for many organisms such as phototrophs whose lifestyle is tailored around light availability and diurnal cycle (Andreas Angermayr et al., 2016; Diamond et al., 2015). To account for metabolite level fluctuations within FBA at a given time point, metabolite accumulation/depletion terms (i.e., the right-hand side (RHS) of mass balance constraints) need to be temporally decoupled. Of course, over time all imbalances will have to average out for a stable culture. In a previous study by Reznik and coworkers, allowing for metabolite imbalances was necessary for the FBA model to consistently capture observed changes in pools of growth-limiting metabolites (Reznik et al., 2013). In another study, relaxing metabolite balance constraints yielded a better correlation between FBA predictions and experimentally observed intracellular metabolic fluxes (MacGillivray et al., 2017). In both studies imbalances for different metabolites were assumed to be independent from one another. However, even under unsteady-state conditions conserved metabolite pools (CM-pools) (Famili and Palsson, 2003) must remain invariant. These pools arise because the overall amount of a particular chemical moiety (such as acyl carrier protein, NAD+/NADH, or folate species) cannot change (in the absence of active biosynthesis) even under the action of the available (exchange) reactions in the system. The mathematical consequence of a CM-pool is that a weighted sum of the RHS terms for metabolites within the pool must equal zero, consistent with a balance between all accumulation (positive RHS) and depletion (negative RHS) terms (Cornish-Bowden and Hofmeyr, 2002; Nikolaev et al., 2005). Thus, any investigation of temporal variation in metabolite levels and resulting phenotypic effects must directly (or indirectly) account for relationships embedded within CM-pools. In the past, CM-pools have been found by either examining the structure of the stoichiometric matrix directly (Cornish-Bowden and Hofmeyr, 2002; De Martino et al., 2014; Famili and Palsson, 2003; Reder, 1988) or examining the feasible region of optimization problems with stoichiometric restrictions included (Nikolaev et al., 2005). As many as 74 aggregated CM-pools, whose subsets of members constitute smaller CM-pools, have been identified for the *Escherichia coli* genome-scale model *i*AF1260 (De Martino et al., 2014).

In this study, we first quantify how uncertainty in the biomass constituent coefficients propagates to FBA results for maximum theoretical biomass yield calculations using the latest *E. coli* model *i*ML1515 (Monk et al., 2017). Two cases were considered: (i) naively ignoring and (ii) considering the need to re-normalize coefficients so that the biomass MW is strictly 1 g mmol^−1^. Failing to rescale biomass reaction coefficients resulted in three-fold higher standard deviation (SD) for the biomass yield upon sampling. Uncertainty in the ATP maintenance parameters had an even greater impact on the predicted biomass yield SD (i.e., six-fold higher than that of biomass composition uncertainty). We found, that in general, any injected random noise (i.e., uncertainty) on biomass coefficients is dampened in the FBA prediction for the biomass yield. This observation is consistent with the surprising robustness of FBA predictions despite the many assumptions inherent in the biomass reaction construction and parameterization. However, predicted intracellular metabolic fluxes can be quite sensitive to the injected random noise. We also explored the effect of allowing for deviations from the metabolite steady-state assumption by temporally relaxing the equality of production and consumption terms (i.e., RHS of balance constraints). We found that FBA problems with randomly sampled non-zero RHS vectors were (almost) always infeasible because CM-pool constraints were violated. Projecting all sampled RHS vectors onto the subspace defined by the CM-pool constraints (thereby ensuring their feasibility) results in a seemingly feasible FBA problem but unless elemental (i.e., C, N, P, etc.) balance constraints are explicitly included, undetected elemental balance violations occur. In steady-state FBA, reaction stoichiometries included in the model are elementally balanced, thus no additional elemental balance constraints are needed. However, whenever the RHS terms are perturbed away from zero, the elemental balances are not automatically satisfied thus requiring their explicit inclusion in any unsteady-state FBA calculations. Additionally, we found that the average FBA predicted biomass yield is significantly decreased compared to the steady-state case. This alludes to a strong selection pressure for strictly maintaining steady-state in metabolite balances, as any departures tend to handicap the cell with a lower (on average) biomass yield potential. This further bolsters the underlying assumption of steady-state in FBA and helps explain its effectiveness in predicting overall cellular phenotypes.

## 2. Methods

### 2.1. Sampling protocol of perturbations in the biomass coefficients and ATP maintenance

We used the latest *E. coli* model (*i*ML1515) (Monk et al., 2017) in minimal media (M9) with glucose as carbon source to determine the effect of uncertainty in biomass coefficients on flux predictions. The engineered reverse beta-oxidation pathway, which is absent in the wild-type strain but included in the model, was turned off by setting the upper bounds of the reactions ACACT1r-8r to zero. The lower bound of pyruvate:flavodoxin oxidoreductase (POR5) was also fixed to zero, as its activity in the reverse direction is only induced by superoxide generators (Nakayama et al., 2013). Model parameters subjected to uncertainty include biomass precursor coefficients, macromolecular compositions, GAM, and NGAM. Parameters were sampled 10,000 times from a normal distribution with the mean centered at the original value and relative SD of 5%, 10%, 20%, and 30% of the original value (hereafter referred to as uncertainty level). During sampling, we ensured that the following material balances were maintained in the biomass reaction: (i) ATP hydrolysis recapitulating GAM, (ii) pyrophosphate in RNA and DNA synthesis, and (iii) water in protein synthesis. Failure to recognize the ATP hydrolysis balance results in a mismatch in ATP and ADP coefficients resulting in either (i) ATP coefficients being higher and its *de novo* synthesis competing with organism growth, or (ii) ADP coefficients being higher creating a metabolic surplus.

Biomass precursor coefficients and the GAM value were first extracted from the biomass reaction with lumped coefficients. Then, random sampling of precursor coefficients or the GAM value was performed. Before reassembling the biomass reaction from sampled parameters, re-normalization of the biomass MW to 1 g mmol^−1^ was performed as:

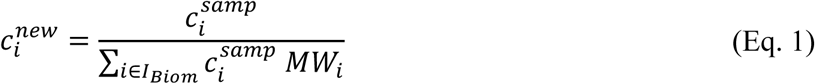

where *I*_*Biom*_ is the set of biomass constituents in the biomass reaction, 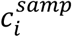 is the randomly sampled coefficient for metabolite *i, MW*_*i*_ is the MW of metabolite *i*, and 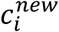 is the recalibrated biomass coefficient for metabolite *i*. For macromolecular composition perturbation, the weight fractions of macromolecular classes were sampled and coefficients of constituents in each class uniformly scaled. Classification of 66 biomass constituents is provided in Supplementary Materials 1 for all amino acids, cell wall components, lipids, RNAs, DNAs, inorganic ions, and cofactors and prosthetic groups.

### 2.2. Quantifying uncertainty propagation from biomass coefficients and ATP maintenance to biomass yield and predicted metabolic fluxes using the standard deviation ratio (SDR) metric

The sampling workflow and propagation of uncertainty is illustrated in Figure 1. For every sample, a biomass reaction was reconstructed and parsimonious flux balance analysis (pFBA) (Lewis et al., 2010) was performed to obtain a predicted flux distribution. Maximum glucose uptake rate was set at 10 mmol gDW^−1^ h^−1^. Mean and SD were calculated for the fluxes of every reaction *j* in the population, and the propagation of uncertainty was quantified using the SD ratio (SDR) defined as:

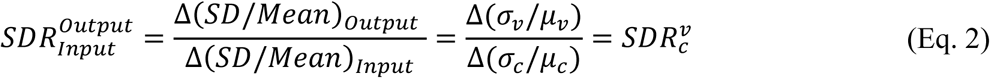

where *σ*_*v*_ and *μ*_*v*_ are the SD and mean for the pFBA output flux or yield *ν, σ*_*c*_ and *μ*_*c*_ are the SD and mean for the input parameter *c* subjected to the injection of normally distributed random noise, and 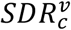 is the rate of change of the output relative SD with respect to the input relative SD. For example, 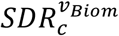 and 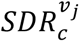 quantify the amplification (or dampening) of the effect of injecting noise on a parameter *c* on the pFBA predicted biomass yield *ν*_*Biom*_ or metabolic flux *ν*_*j*_, respectively. SDR was calculated with linear regression for the calculated output over the input SD.

**Figure 1.**
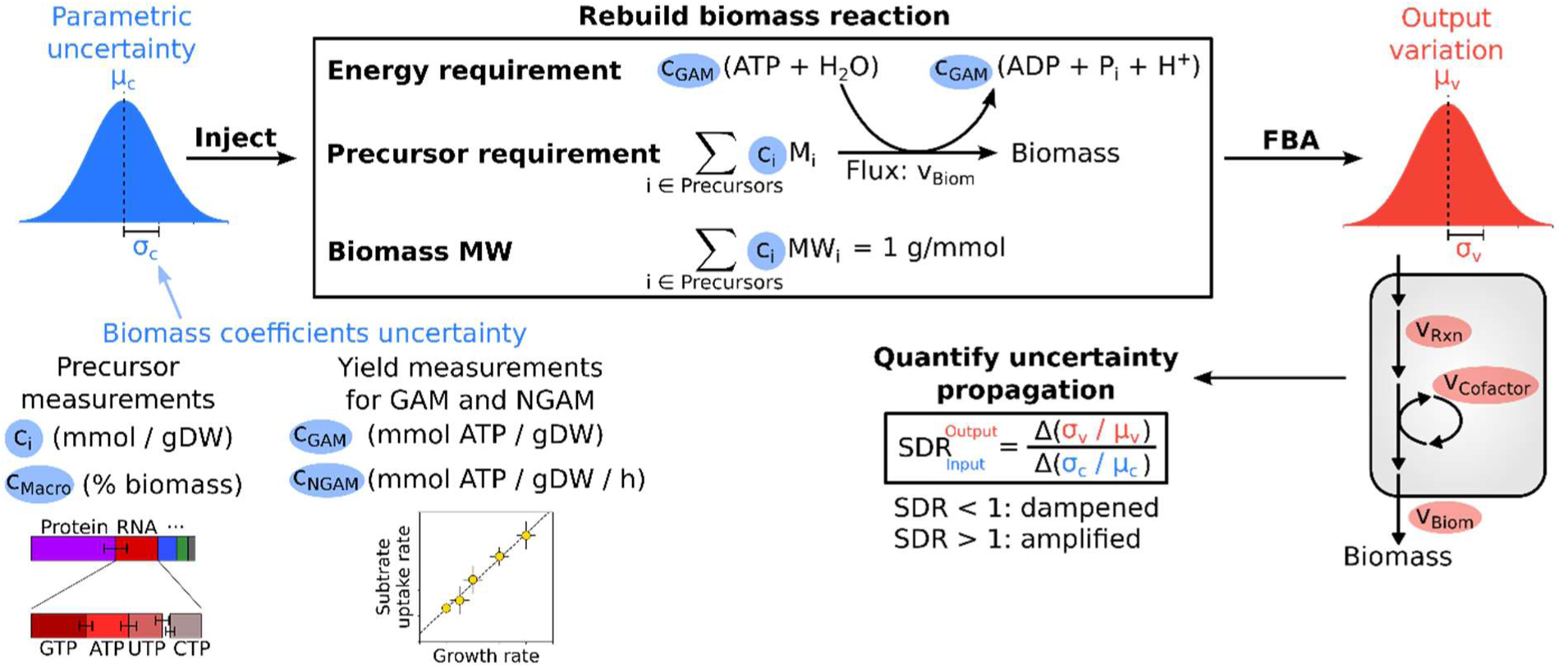
Overview of the method to inject and quantify propagated uncertainty from biomass coefficients and ATP maintenance rates to FBA predictions. First, normally distributed uncertainty is injected to biomass precursor coefficients and ATP maintenance parameters. The biomass reaction is reassembled while accounting for ATP hydrolysis balance and biomass MW. FBA is next carried out using the new biomass reaction and parameter 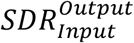 quantifies the extent of uncertainty propagation.

NADH, NADPH, and ATP cofactor production flux SDRs 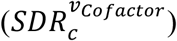 were calculated by aggregating over all cofactor producing reactions (*ν*_*Cofactor*_),

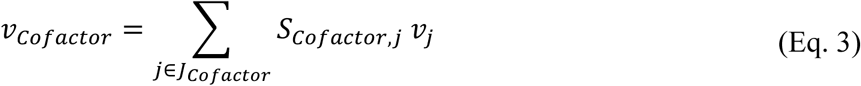

where *J*_*Cofactor*_ is the set of reactions producing a particular cofactor, *S*_*Cofactor,j*_ is the cofactor stoichiometry in reaction *j*, and *ν*_*j*_ is the flux through reaction *j*. See Supplementary Materials 1 for a list of all ATP, NADPH, and NADH producing reactions.

SDR for the sum of fluxes through alternative (or parallel) pathways was computed in a similar manner to Eq. 2, by summing all fluxes connecting the same start and end metabolites. This is done to quantify the uncertainty propagating to the net conversion flux, which is typically more dampened than to the individually distributed flux components through alternative pathways. See Supplementary Materials 1 for a list of all “flux sum” quantities and the corresponding reaction sets.

### 2.3. Protocol for sampling departures from the metabolite steady-state and instationary FBA

Departure from metabolite steady-state means one or more RHS terms of the FBA’s mass balance constraints (Orth et al., 2010) (denoted by *b*_*i*_ for metabolite *i*’s balance (Nikolaev et al., 2005)) may assume non-zero values. A positive (or negative) RHS term indicates metabolite accumulation (or depletion). Randomly sampled RHS terms (almost) always lead to a setup of an infeasible optimization FBA problem because CM-pool constraints are consistently violated. These include CM-pool constraints for a subset of RHS terms corresponding to the conservation of chemical moieties (e.g., ACP or CoA) in metabolites (De Martino et al., 2014; Nikolaev et al., 2005; Reder, 1988) that cannot be degraded (and *de novo* synthesized for some moieties) due to a lack of degradation (and biosynthesis) pathway in the model (Chan et al., 2017). In addition, some nutrient metabolites and their derivatives form a CM-pool in a specific growth condition (but not in others) if the corresponding exchange reactions that supply the required nutrients are blocked. Thus, for metabolites in CM-pools, their accumulation is coupled with the depletion of other metabolites. As the first step in identifying these coupled metabolites, we introduce an optimization formulation called <u>c</u>onserved <u>m</u>etabolite <u>p</u>ool <u>check</u> (*CMP-check*) that checks whether a given metabolite *i*^∗^ is part of a CM-pool. The key idea here is to introduce an imbalance *ε* denoting accumulation of metabolite *i*^∗^ (or −*ε* for depletion), and then determine if (i) it is coupled to steady-state perturbations to other network metabolites, (ii) can be satisfied through exchange with the growth medium, or (iii) is always infeasible. Slack variables 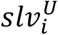 and 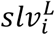 denote positive and negative departures, respectively, from the steady-state condition for metabolite *i* (*i* ≠ *i*^∗^). The objective of *CMP-check* is the minimization of the sum of slack variables across all metabolites, thus identifying the minimal set of coupled (imbalanced) metabolites. The formulation is as follows:

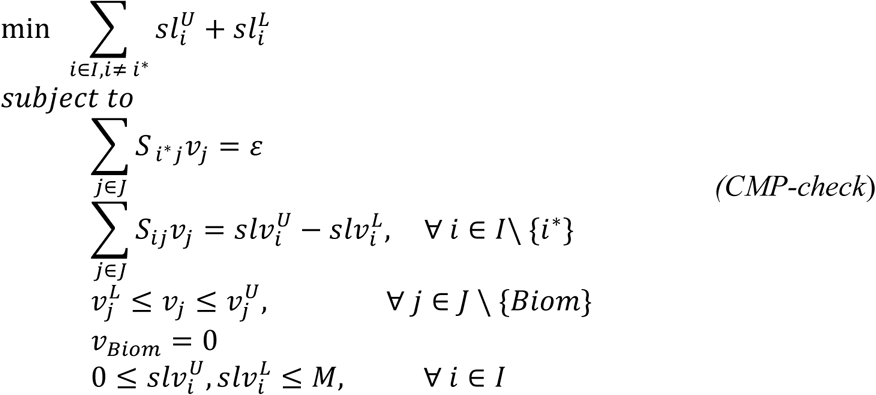

where *M* is an arbitrarily large scalar. Flux through the biomass reaction (*ν*_*Biom*_) was turned off to prevent metabolite rebalancing through accumulation (or depletion) of the biomass sink flux. *CMP-check* was solved twice for every metabolite *i*^∗^ for both accumulation (using *ε*) and depletion (using −*ε*). We used an arbitrarily small value of 0.01 mmol gDW^−1^ h^−1^ for *ε*, and 1,000 mmol gDW^−1^ h^−1^ for *M*.

Because generating of CM-pool specific constraints is computationally inefficient for genome-scale models (see Supplementary Text), we hereby introduce a workflow that injects uncertainty into metabolites’ RHS values and quantifies uncertainty propagating to FBA-predicted biomass yield while accounting for CM-pools and elemental balance constraints. Imbalances were introduced to the RHS terms of the metabolite mass balance constraints (i.e., by an amount 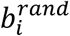) by sampling from a normal distribution centered at zero at the uncertainty levels *σ*^*b*^ of 0.1%, 1%, and 10% of the glucose uptake rate by mass (i.e., in the unit of gram glucose gDW^−1^ h^−1^). We start from the mass-basis SD *σ*^*b*^ (i.e., unit of gram gDW^−1^ h^−1^) to avoid the injecting of larger deviation to metabolites with larger MW values. For example, molar-basis sampling of the accumulation rate of the metabolite “(O16 antigen) x4 core oligosaccharide lipidA” (o16a4colipa_p), whose MW is 44-fold higher than that of glucose, will introduce a disproportional drain of resource in the system. SD in the molar unit (i.e., mmol gDW^−1^ h^−1^) (i.e., denoted as 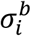) consistent with the unit of the RHS term can be calculated using the following equation:

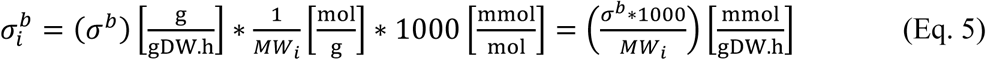

where *MW*_*i*_ is the MW of metabolite *i*. Using optimization formulation *CMP-check*, metabolites were split into three groups: (i) allowed to only accumulate (i.e., 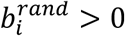), (ii) allowed to only go into deficit (i.e., 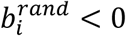), and (iii) allowed to either accumulate or go into deficit (i.e., 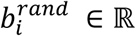). Steady-state departure variables 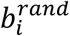 were then sampled from a half-normal distribution for cases (i) and (ii) and from a full normal distribution for case (iii). The normal distribution was centered at zero with a SD of 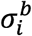. A half-normal distribution is equivalent to a chi-square distribution with a single degree-of-freedom and scaled by 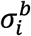.

To obtain a vector ***b***^***feas***^ composed of individual RHS terms 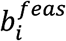 for FBA, we introduced and used the Projection Using Linear Programming (*PULP*) formulation that projects the vector ***b***^***rand***^ composed of individual RHS terms 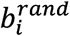 to the subspace that satisfies network stoichiometry and reaction directionality restrictions:

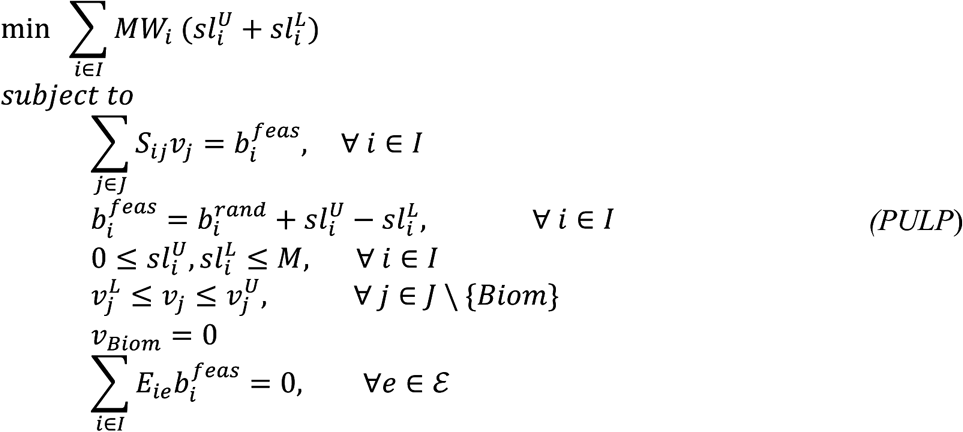

where 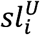 and 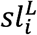 are positive and negative slack variables, respectively, and *E*_*ie*_ denotes the stoichiometry of element or moiety *e* in metabolite *i*’s chemical formula. The set of all elements and moieties collected from the metabolites’ formulae is denoted by *ε*. By minimizing the sum of slack variables 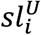 and 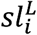, the initial randomly generated RHS vector ***b***^***rand***^ is thus projected onto the feasible RHS solution space. The last constraint in *PULP* denotes elemental balance constraints dictating the conservation of elements, such as C, N, and P.

10,000 RHS vectors (***b***^***rand***^) were randomly sampled, as described above, for each uncertainty level *σ*^*b*^ and corresponding feasible RHS vectors (***b***^***feas***^) generated using *PULP*. In order to quantify the effect of metabolite concentration departures from the steady-state condition, we introduce the instationary *FBA* (*iFBA*) version of FBA where the RHS terms *b*_*i*_ of mass balance constraints are set to 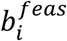 to yield a feasible optimization problem. The optimization formulation for a standard growth-maximization *iFBA* is formulated as:

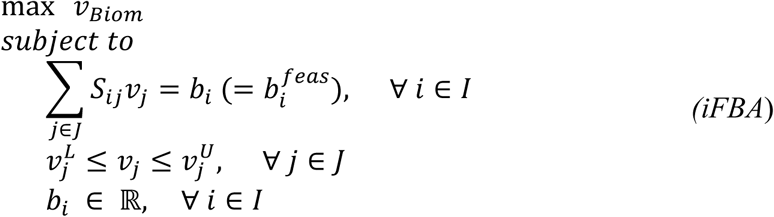

where *I* and *J* are the set of metabolites and reactions, respectively, *S*_*ij*_ is the stoichiometric coefficient of metabolite *i* in reaction *j, ν*_*j*_ is the flux of reaction *j, ν*_*Biom*_ is the specific growth rate or biomass dilution flux, and 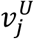 and 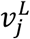 are the upper and lower bounds of reaction *j*, respectively. 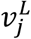 assumes a value of zero for irreversible reactions and an arbitrarily large negative value for reversible reactions. Then, the mean and SD of *ν*_*Biom*_ normalized by steady-state FBA’s *ν*_*Biom*_ are calculated to quantify the output variation.

Note that in the absence of elemental balance constraints, the RHS vector generated by *PULP* (almost) always introduces a net deficit or surplus of chemical elements to the system. We also run the *PULP* without the elemental balance constraints yielding elementally imbalanced ***b***^***feas***^ for which the elemental surplus for an element *e* normalized by the glucose uptake rate by mass can be quantified as:

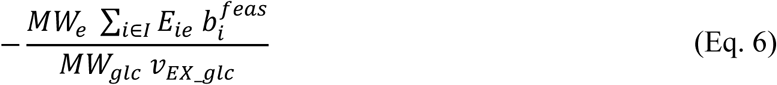

where *MW*_*e*_ is the atomic weight for the element *e, MW*_*glc*_ is the molecular weight of glucose, and *ν*_*EX_glc*_ is the glucose uptake rate. A positive elemental surplus value means more moles of an element are consumed than produced leading to a metabolic surplus, and vice-versa for a negative value.

Because metabolites’ formulae are used to derive the elemental balance constraints, the identity of all generic “R” and “X” groups were disambiguated into 16 specific conserved moieties using the metabolite names for 278 metabolite formulae in the *i*ML1515 model. For metabolites’ MW used throughout the analysis, a zero value was arbitrarily assigned for the MW of the conserved moieties (i.e., formerly “R” and “X” groups) since they are not provided with the model. Thus, the MW of metabolites with a partially defined formulae, such as acyl carrier protein (i.e., formula C_11_H_21_N_2_O_7_PRS) is calculated after omitting undefined groups. Updated metabolite formulae and their MW values are provided in the Supplementary Materials 2.

### 2.4. Software implementations

Sampling of biomass coefficients was carried out using the Python 3.6 package NumPy for normal distribution sampling (numpy.random.normal) (Virtanen et al., 2020) and COBRApy for flux balance analysis (Ebrahim et al., 2013). Sampling of metabolite RHS terms was carried out using the Python 3.6 package SciPy for normal and half-normal distribution sampling (scipy.stats.norm and scipy.stats.halfnorm) (Virtanen et al., 2020). *CMP-check* and *PULP* were implemented in General Algebraic Modeling System (GAMS) programming language (version 33.2.0, GAMS Development Corporation), with IBM ILOG CPLEX (version 12.10.0.0) as the solver. Example scripts are available at https://github.com/maranasgroup/uncFBA.

## 3. Results

We first describe how the injection of normally distributed random noise into biomass coefficients and ATP demand propagates in FBA predictions towards biomass and metabolic flux predictions, and assess the sensitivity (or lack thereof) of predicted flux distributions to input parameters.

### 3.1. Propagation of uncertainty from biomass coefficients and ATP maintenance to FBA predictions of metabolic fluxes and biomass yield

The overall workflow which is described in detail in the Methods section (Section 2.1 and 2.2) is illustrated using Figure 1. We quantify how biomass coefficients and/or ATP maintenance uncertainty is propagated to FBA predicted biomass yield and metabolic fluxes.

In all cases, the effect of injecting random noise to the biomass coefficients is dampened in the biomass yield predictions by FBA. This is because the random introduction of noise causes some of the coefficients to increase but others to decrease, thus partially counteracting any changes in the overall demand for biomass formation. This is a consequence of the fact that the biomass flux is affected by many constituent coefficients and unless they all change in a coordinated manner, the effect on the biomass flux is attenuated. Even if some random noise injections happen to cause a coordinated increase in the demand for some constituents for the formation of one unit of biomass, perturbations for other constituents are likely to cancel them out. Furthermore, as seen in Figure 2A, by renormalizing the biomass coefficients such that biomass MW is maintained at 1 g mmol^−1^, the propagation of uncertainty to the biomass yield is revealed to be even smaller than by naively ignoring this renormalization. The relative magnitude of the noise amplitude decreases from 14.9% to 4.9% as quantified by the standard deviation ratio 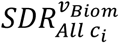 metric (Eq. 2). Uncertainty in the macromolecular composition leads to a higher 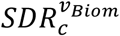 (i.e., 7.1% vs. 4.9%, see Figure 2B). Note that in a previous study examining biomass coefficient uncertainty (MacGillivray et al., 2017), the effect of omitting MW-normalization was not detectable because the imposed uncertainty was much smaller (i.e., between ∼10^−1^ to 10^−2^ <u>± 10^−6^</u> mmol gDW^−1^ for most metabolites) as compared to this study (i.e., between ∼10^−1^ to 10^−2^ mmol gDW^−1^ <u>± 10 to 30%</u>).

**Figure 2.**
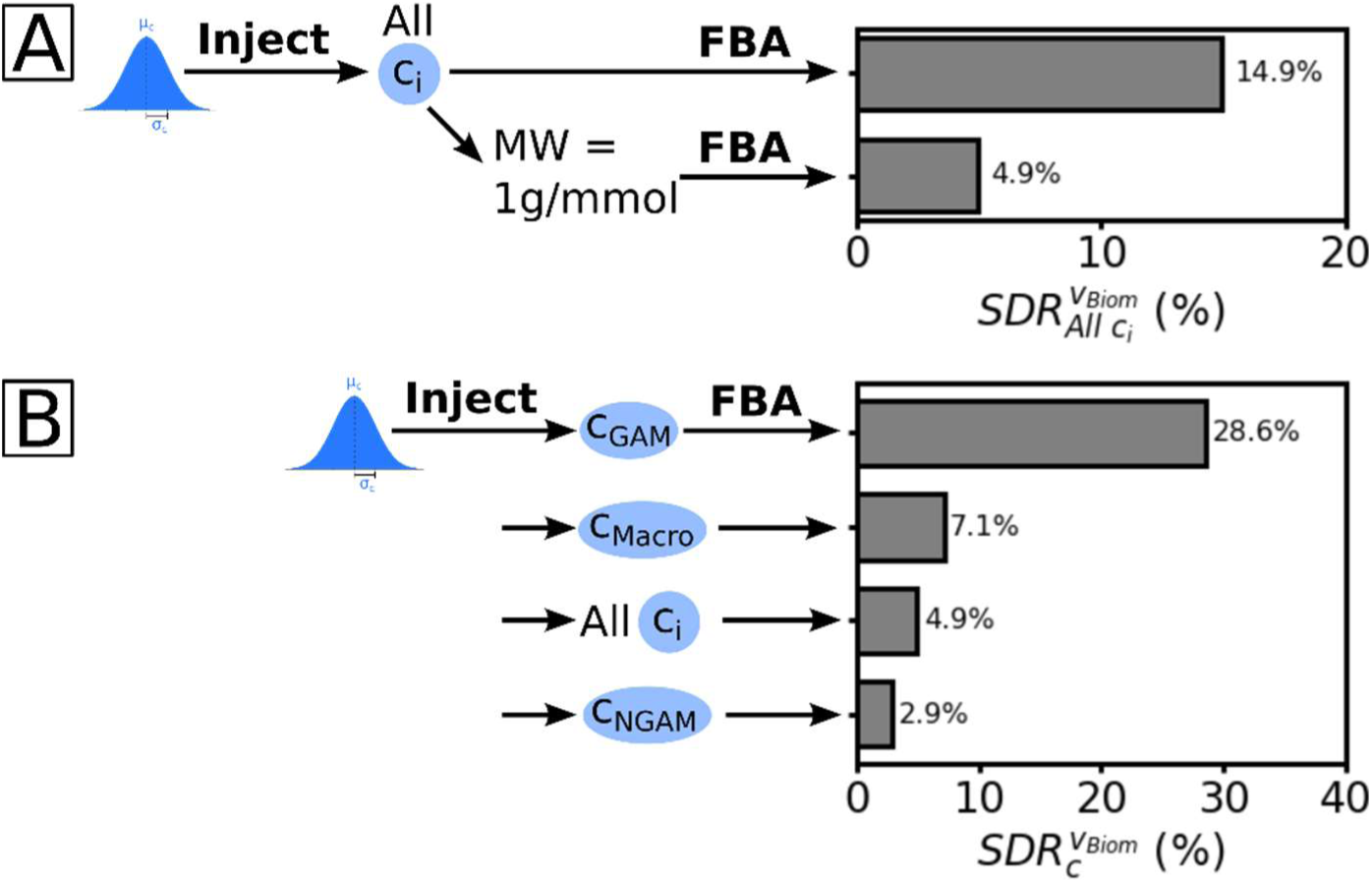
Effect of uncertainty in biomass composition and ATP demand on FBA predictions. (A) Neglecting to reweigh biomass MW to 1 g mmol^−1^ leads to a 3-fold exaggerated effect to the biomass yield output. (B) Contribution of individual biomass coefficients (*c*_*i*_), macromolecular composition (*c*_*Macro*_), GAM (*c*_*GAM*_), and NGAM (*c*_*NGAM*_) uncertainty on biomass yield output.

SDR metrics for the biomass yield and cofactor net production fluxes are provided in Table 1. We find that random noise propagation is highly dampened (i.e., SDR < 10% with few exceptions) for both biomass yield and cofactor fluxes. This may explain why FBA is surprisingly successful at predicting biomass and/or flux yields despite the highly uncertain and/or variable nature of required inputs to describe the carbon and energy tally for biomass formation. Because NADPH is the primary redox cofactor for biosynthesis of biomass constituents, FBA predicted variance for the total NADPH synthesis flux (i.e., 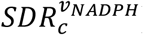) under biomass composition uncertainty is higher than those for biomass, NADH or ATP (see Table 1).

**Table 1.** Calculated SDR values for biomass yield and cofactor net production due to propagated uncertainty from biomass coefficients and ATP maintenance parameters. Do note that biomass MW is reweighted to 1 g mmol^−1^ unless mentioned otherwise in the remainder of the manuscript.

| Sampled parameters | $SDR_c^{v_{Biom}}$ | $SDR_c^{v_{NADPH}}$ | $SDR_c^{v_{NADH}}$ | $SDR_c^{v_{ATP}}$ |
| --- | --- | --- | --- | --- |
| *Species composition* |  |  |  |  |
| All coefficients | 4.9 | 10.1 | 5.5 | 5.9 |
| All coefficients (unnormalized MW) | 14.9 | 11.2 | 8.9 | 10.6 |
| *Macromolecular composition\** |  |  |  |  |
| All compositions | 7.1 | 16.3 | 9.6 | 9.3 |
| RNA fraction (22%) | 5.0 | 8.2 | 6.4 | 7.0 |
| Lipid fraction (10%) | 4.2 | 0.3 | 0.8 | 3.2 |
| Protein fraction (58%) | 1.8 | 13.9 | 7.2 | 5.1 |
| Inorganic ions fraction (1%) | 0.7 | 0.3 | 0.4 | 1.3 |
| Cell wall fraction (6%) | 0.5 | 2.5 | 0.3 | 1.1 |
| DNA fraction (3%) | 0.5 | 1.2 | 0.7 | 0.8 |
| Cofactors fraction (0.3%) | 0.01 | 0.1 | 0.1 | 0.7 |
| *ATP maintenance* |  |  |  |  |
| Growth-associated (GAM) | 28.60 | 28.48 | 38.73 | 46.81 |
| Non-growth associated (NGAM) | 2.92 | 2.93 | 3.94 | 4.94 |

\*Number in brackets is the mass fraction of the respective macromolecule class in the *iML1515 E. coli* GSM's biomass reaction [10].

In contrast to imposing random noise on all biomass constituent coefficients at the same time, change in only the GAM requirement causes more pronounced changes in the biomass yield and cofactor balances (see Table 1 and Figure 2A). This is expected as only a single coefficient for GAM undergoes a fluctuation causing systemic changes in ATP demand. This offers a cautionary tale, as a common practice in model reconstruction is to port unchanged ATP maintenance values from previously built models for related organisms even though GAM values are highly variable (e.g., yeasts: 23 – 167 (Aung et al., 2013; Dinh et al., 2019; Mishra et al., 2018; Tomàs-Gamisans et al., 2018, 2016; Torres et al., 2019) & bacteria: 18 – 75 mmol gDW^−1^ h^−1^ (Flahaut et al., 2013; Kavvas et al., 2018; Liao et al., 2011; Monk et al., 2017; Orth et al., 2011; Seif et al., 2019; Thiele et al., 2011)). Growth conditions such as supplied substrate (Tomàs-Gamisans et al., 2018, 2016) and nutrient- and oxygen-limited growth (Dinh et al., 2019; Torres et al., 2019) can also cause significant departures. In contrast, changing only the coefficient for NGAM caused a much smaller change in the predicted biomass yield (i.e., 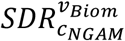 of 2.9%). This is a reflection of the relative demands imposed by GAM and NGAM, respectively, on the consumed glucose (i.e., 27% and 3%). The effect of injecting noise on precursor demands for biomass formation (i.e., coefficients or macromolecular compositions) was significantly smaller even though a much larger fraction of glucose (i.e., ∼71%) is allocated to their synthesis. Again, the reason for this insensitivity is that the injection of random noise tends to alter their relative proportions but not the total mass (i.e., 1 gram of precursors to make 1 gram of biomass). In contrast, random noise on GAM or NGAM requirements directly increases or decreases the net ATP demand without a compensating mechanism. This implies that random errors for coefficients of multiple biomass constituents in a genome-scale model are likely to lead to an attenuated response of FBA biomass yield predictions, whereas systematic errors that directly affect the overall carbon and energy demand will likely cause changes in FBA prediction in proportion to the magnitude of the error.

Zooming in closer we also examined the sensitivity of the biomass yield upon injecting noise one-at-a-time to macromolecular groups (see Table 1 and Figure 2C). As expected, uncertainty in macromolecules accounting for a larger fraction of biomass (by mass) gave rise to a higher 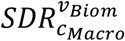 value. These include proteins (58% by weight), RNAs (22% by weight), and lipids (10% by weight). The relative order is determined not only by the net weight fraction, but also by the amount of carbon, ATP, and NAD(P)H consumed during biosynthesis. Even though RNAs and lipids rank after proteins in terms of contributing biomass weight fractions, their uncertainty propagation has a larger effect in biomass yield (Table 1). This is due to higher cofactor costs when forming more covalent bonds for larger RNA (C_9_ – C_10_) and lipid (C_37_) molecules compared to amino acids (C_2_-C_9_). Interestingly the effect on NAD(P)H production upon the introduction of random noise in the lipid fraction is much less than the effect on biomass yield (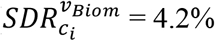 vs. 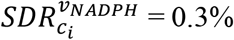 and 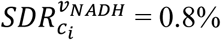). This is due to cofactor promiscuity in enoyl-ACP reductase (catalyzing the last step in a fatty acid elongation cycle) (Bergler et al., 1996) allowing the use of NADH in place of NADPH as the reducing equivalent. As a result, equimolar consumption of ATP, NADH, and NADPH occurs per fatty acid elongation cycle (out of 16 cycles in total). The large but almost equimolar total requirement of 18 ATP, 17 NADPH, and 16 NADH in mol/mol for lipid synthesis could explain the reason for its effect on biomass yield but not cofactor availability as the equimolar cofactor requirement allows the cofactor load to be distributed across different cofactor producing parts of the network. In contrast, cofactor demand of protein synthesis is skewed towards NADPH (e.g., L-methionine requires 8 NADPH, 5 ATP, and 1 NADH in mol/mol) whereas cofactor demand of RNA synthesis is skewed towards more ATP (e.g., *de novo* synthesized ATP requires 6 NADPH and 11 ATP in mol/mol), creating disproportionally heavier loads to NADPH and ATP, respectively which is reflected in the calculated SDR (Table 1). This observation is consistent with the experimentally observed importance of controlling NADPH metabolism in amino acid producing strains of *E. coli* (Xu et al., 2018). Lastly, while minor fraction components such as cofactors and inorganic ions are important to include in expanding the resolution of FBA predictions (Feist and Palsson, 2010), the effect of introducing noise in their respective biomass coefficients on biomass yield is negligible as they constitute only a small fraction (i.e., ∼1% of biomass by weight, all 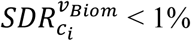).

Next, we examined the effect of injecting random noise (one-at-a-time) to biomass coefficients and quantified the output SDR (see Figure 3A and Supplementary Material 1) to determine biomass precursors that have a disproportionally large effect on FBA predictions. As seen previously, uncertainty propagation partially correlates with how much each precursor contributes to biomass by mass (i.e., gram per gram biomass, %) (see Figure 3B). Exceptions to this general trend include the biomass constituents of two phosphoethanolamines (i.e., lipid species) affecting biomass yield, GTP and ATP (i.e., RNA species) affecting ATP production, and L-methionine and L-isoleucine (i.e., amino acids) affecting NADPH production due to their differences in cofactor costs, as explained for macromolecular fraction uncertainty. GTP and ATP biomass coefficient values also affect NADH production, as NADH fuels oxidative phosphorylation to generate ATP. Interestingly, NADH production is affected by the L-leucine biomass coefficient because it is produced in the leucine biosynthesis pathway: one from 3-isopropylmalate dehydrogenase and one from producing the precursor acetyl-CoA (from pyruvate via pyruvate dehydrogenase). Based on the reported SDR values (see Supplementary Material 1) this analysis can be used to prioritize measurements of biomass components that could improve the accuracy of flux and yield quantification.

**Figure 3.**
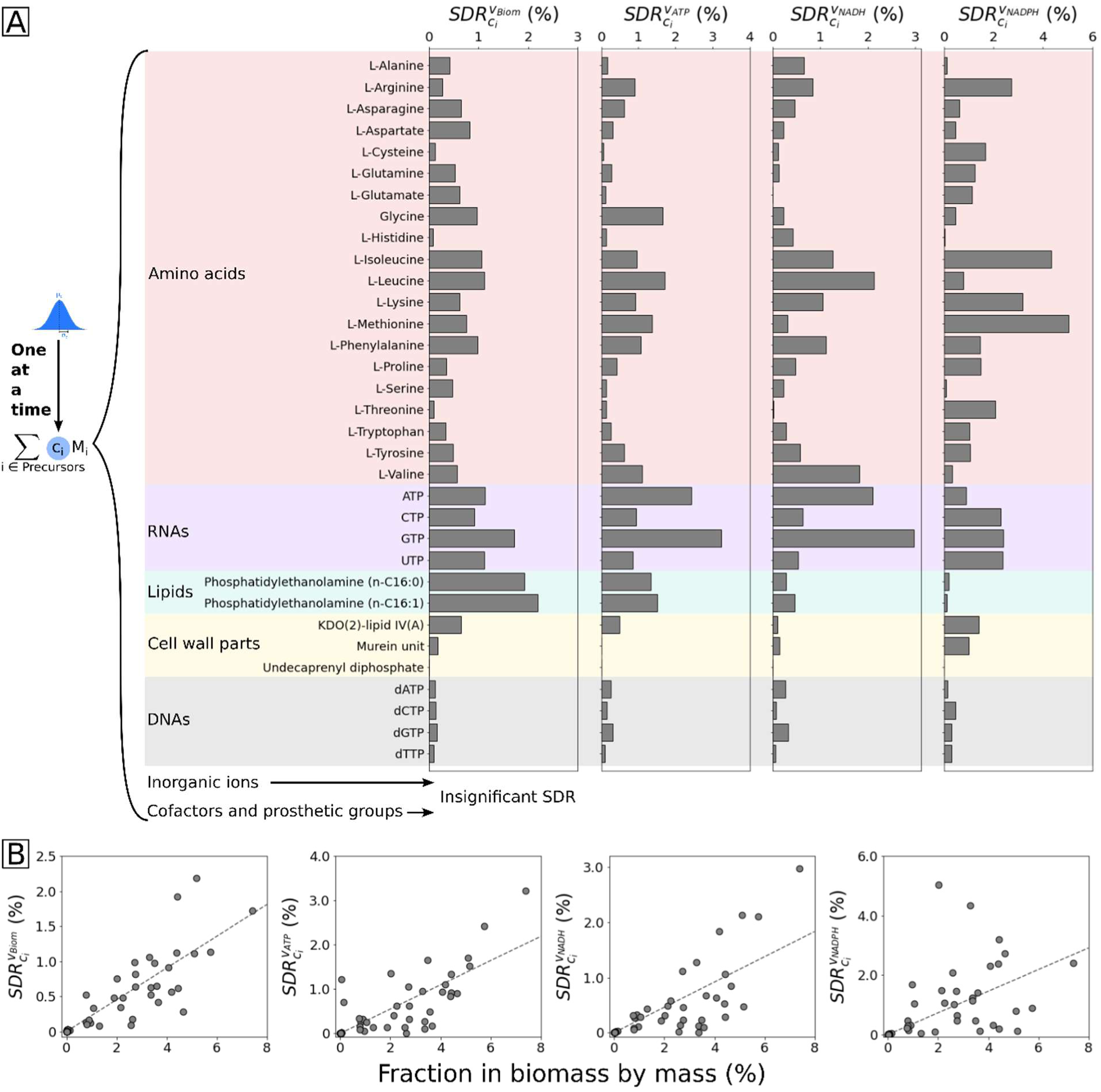
(A) Responses in output standard deviation for biomass yield and cofactor fluxes due to “one-at-a-time” noise injection to biomass coefficients. From left to right are responses in the output standard deviation ratios of biomass yield, ATP, NADH, and NADPH fluxes. (B) Plots of the aforementioned SDR values vs. coefficient mass fraction by mass in biomass composition. The dotted lines are the linear regression lines.

Unlike biomass yield or aggregated demands for cofactors, injecting random noise in biomass coefficients can cause varied, and in some cases, an amplified effect for some metabolic fluxes and pathways (see Figure 4). Central metabolism fluxes exhibited higher SDR values under GAM uncertainty whereas downstream biosynthesis fluxes had larger SDR under biomass composition uncertainty. Notably the SDR for the pentose phosphate pathway was consistently high under either GAM or biomass coefficients uncertainty. Under NGAM uncertainty, as expected, all reaction 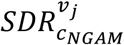 values were below 25% (median = 2.9%). Even though on average the uncertainty propagation from biomass composition random noise injection is still less than proportional for most of the fluxes, it is significantly larger (see Figure 4) than the one for biomass yield (see Table 1). This implies that biomass composition accuracy is more important for predicting individual metabolic fluxes using FBA as opposed to the overall biomass yield.

**Figure 4.**
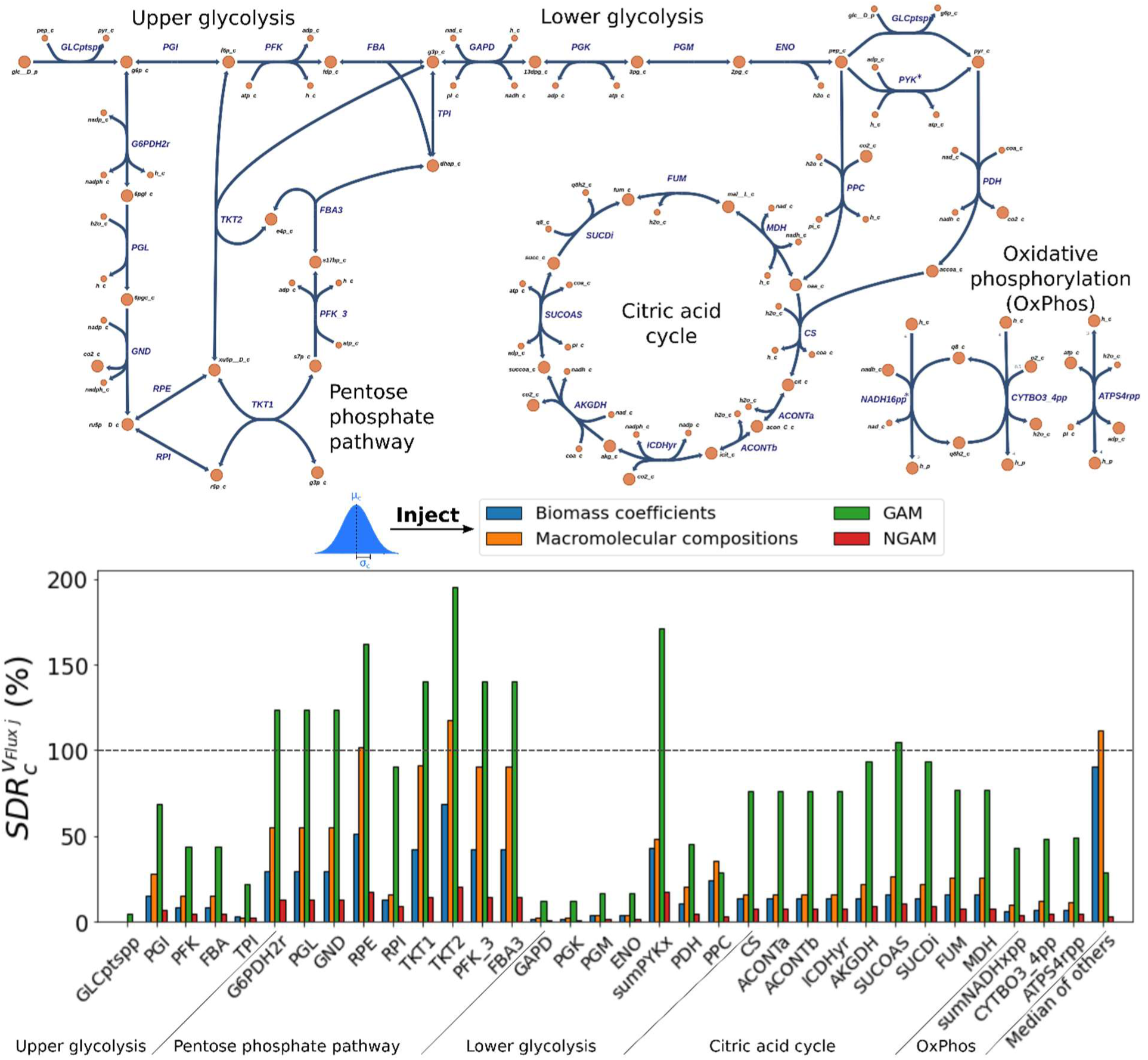
SDR of individual fluxes in central metabolism. Reaction IDs used in the metabolic map and the bar chart follow the BiGG notation used by the model *i*ML1515 [31]. The last entry, “median of others”, corresponds to the median SDR for all reactions that are not part of central metabolism (i.e., mostly biosynthesis reactions).

Starting from the extracellular glucose node, the phosphotransferase system (PTS) exhibited the least variation among all reactions upon random noise injection to the parameters (see Figure 4, reaction GLCptspp in “upper glycolysis”). The maximization of biomass yield in FBA indirectly couples it with maximal glucose uptake. Further downstream, glucose 6-phosphate (G6P) is funneled towards either G6P isomerase (PGI, in glycolysis) or dehydrogenase (G6PDH2r, in PPP). We found that the flux towards G6PDH2r exhibited about twice the sensitivity to the injected uncertainty than towards PGI. This is because PGI funnels twice as much flux as G6PDH2r (i.e., 65% vs. 35% of glucose uptake). The split portions rejoin in lower glycolysis, which showed relatively low sensitivity to parametric uncertainty except for the last pyruvate kinase step (Figure 3). This is because pyruvate is a precursor for many biomass constituents as well as acetyl-CoA used in TCA cycle to produce additional energy. After pyruvate, fluxes in energy-producing pathways of TCA cycle and oxidative phosphorylation are relatively sensitive to GAM uncertainty as the net ATP demand on the system is affected. The pentose phosphate pathway (PPP, producing NADPH and various biomass precursors) was also affected by GAM uncertainty, as an increase in energy demand requires more glucose to be utilized to produce ATP, in turn releasing more CO_2_ as the respiratory byproduct. In particular, transketolase-2 reaction (TKT2) exhibited the highest sensitivity possibly because it is the only reaction in the pFBA solution connecting PPP to the fructose-6P (F6P) node, which is a precursor to the murein unit (i.e., a cell wall component) in biomass. Thus, a coordinated increase or decrease in the demand of PPP metabolites and F6P affects TKT2 more than other PPP reactions that are not connected to the F6P node. Beyond glycolysis, a low sensitivity to biomass composition perturbation was observed in the TCA cycle and oxidative phosphorylation, indicating a relatively stable cellular demand for ATP demand despite fluctuations in composition that tend to counteract one another.

Biomass coefficient variability can also affect metabolism by activating pathways that did not carry flux in the pFBA solution using the original biomass coefficients, where a total of 434 reactions carry flux at maximal growth under glucose utilization. Upon injecting noise to biomass coefficients or GAM uncertainty, this number increases to 471 indicating that alternative pathways may become active in response to a perturbed biomass composition or ATP demand (see Supplementary Text for the list of activated alternative pathways). This implies that changes in the value of the coefficients in the biomass objective function in pFBA can update the optimality order of vertices in the polytope describing the phenotypic space. In some cases, activations are trivial such as switching between quinone and menaquinone but in other cases, physiologically relevant pathway activations are observed involving the use of an alternative redox cofactor (e.g., NADH or NADPH) or energy molecule (e.g., ATP or GTP). Other reaction activation cases involve short linear pathways such as the ones in glycine cleavage (GLYCL), some reactions in one carbon metabolism (FTHFD, FORtppi, sumFDHxpp; Figure 5), GTP biosynthesis (sumGTPsyn), and hydrogen peroxide production (SPODM, sumQMOx).

**Figure 5.**
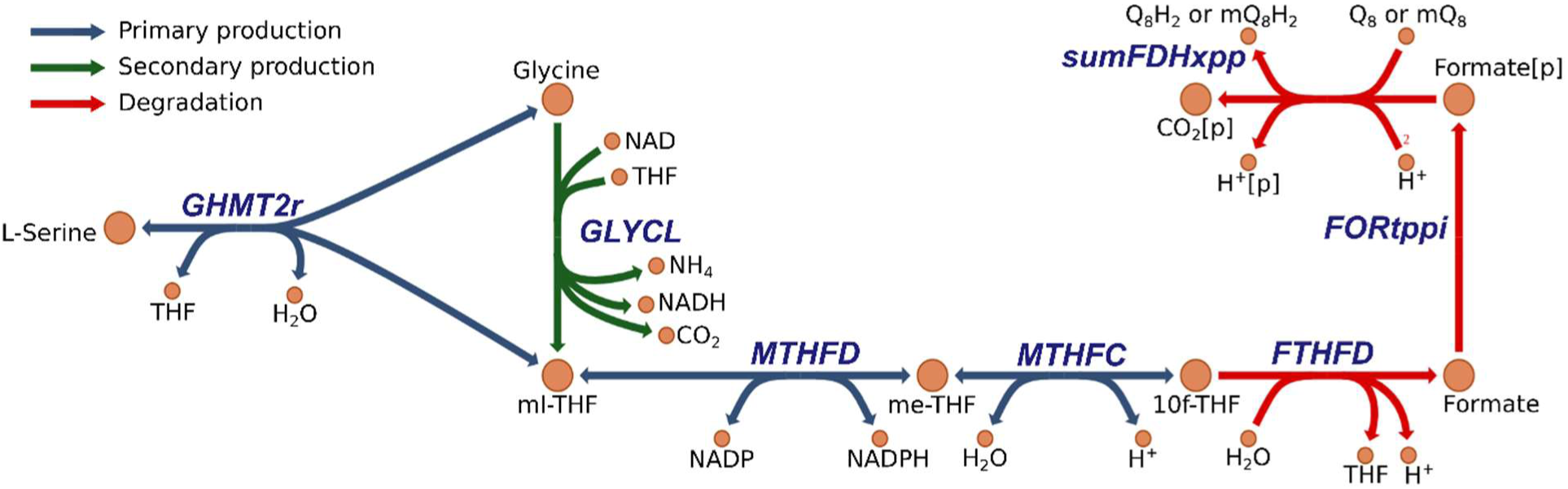
Overview of the one carbon metabolism pathway. GLYCL (highlighted in green) complements 10f-THF production whereas its degradation pathway (highlighted in red) is used to degrade excessive 10f-THF. [p]: in periplasm compartment; THF: tetrahydrofolate; ml-THF: 5,10-methylene-THF; me-THF: 5,10-methenyl-THF; 10f-THF: 10-formyl-THF; Q_8_: ubiquinone-8; Q_8_H_2_: ubiquinol-8; mQ_8_: menaquinone-8; mQ_8_H_2_: menaquinol-8; GHMT2r: glycine hydroxymethyltransferase, GLYCL: glycine cleavage system, MTHFD: methylene-THF dehydrogenase; MTHFC: methenyl-THF cyclohydrolase; FTHFD: formyl-THF deformylase; FORtppi: formate transport via diffusion; sumFDHxpp: formate dehydrogenase (Q_8_ or mQ_8_).

Variability in the coefficients of biomass precursors (such as RNAs) that require one-carbon moiety in the form of 5,10-methylene-tetrahydrofolate (ml-THF) or 10-formyl-tetrahydrofolate (10f-THF) (see Figure 5) can have systemic effects through one carbon metabolism. In *E. coli*, one-carbon moiety is co-produced from glycine via the glycine hydroxymethyltransferase (see Figure 5, “primary production”). In most samples, biomass coefficients for precursors requiring one-carbon moiety are high enough that additional contribution of the moiety from GLYCL is required (see Figure 5, “secondary production”) and only in limited samples (∼6% cases for 10% and ∼30% cases for 30% uncertainty) the moiety was produced in excess. The surplus amount was degraded to formate (via 10f-THF deformylase (FTHFD)) and subsequently to CO_2_ (FORtppi, sumFDHxpp). Interestingly, the activation of this degradation pathway is consistent with the physiological role of FTHFD in controlling the one-carbon pool (Nagy et al., 1995). Another pathway activated under biomass coefficient noise injection was in GTP production (sumGTPsyn), which exhibited a bimodal distribution of flux values across samples. This was due to the presence of two alternative adenylate kinase reactions phosphorylating AMP which can use either ATP (reaction ADK1) or GTP (reaction ADK3) (Rogne et al., 2018). Finally, hydrogen peroxide (H_2_O_2_) production, via quinol monooxidase (sumQMOx) and then superoxide dismutase (SPODM), followed a normal distribution with a relatively wide spread 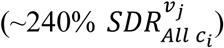. This is because this pathway is not the sole H_2_O_2_ source in the network and is only used in a subset of samples to supplement production of the biomass constituent Fe^3+^ (via reaction FESD1s). Overall, analyzing the activation of pathways upon injection of noise in biomass composition can offer clues as to the optimality criteria that govern the utilization of specific pathways over others and pinpoint which alternatives may become activated when the overall carbon and energy demands are altered.

### 3.2. Propagation of temporally decoupled metabolite balances on FBA biomass yield predictions

Equality of production with consumption for all metabolites is a key assumption that imposes restrictions on FBA outputs. However, metabolism need not obey this stringent criterion at all time-points of the growth cycle, thereby allowing for some surpluses or deficits that can later be evened out. These temporary departures from the strict steady-state assumption may reflect inherent noise in the rate of reactions (Levine and Hwa, 2007), variability among cellular subpopulations in a culture (Kumar et al., 2020), coordinated responses to changes in the microenvironment (Schmitz et al., 2017), or cellular programming (Andreas Angermayr et al., 2016). Sampling unsteady-state regimes may offer the cell an opportunity to jump start metabolism by depleting a stored metabolite pool (i.e., glycogen storage (Shinde et al., 2020)) or deplete oxygen before an oxygen-sensitive enzyme is produced (Milligan et al., 2007; Rabouille et al., 2014). Herein we aim to explore the allowed magnitude of these departures and their overall effect on fitness as quantified by biomass yield. By randomly perturbing the RHS terms of the metabolite balances we (almost) always end up with an infeasible *iFBA* problem (see Section 2.3). This implies that any departure from metabolite steady-state must satisfy the constraints associated with the conservation of conserved metabolite pools (CM-pools, see Section 2.3).

Ignoring these constraints overestimates the permitted variability in the network and results in infeasible solutions. One can attempt to classify all metabolites as to their participation CM-pools by examining one of them at a time (see *CMP-check* formulation in Section 2.3). Three outcomes are possible. A metabolite is part of a CM-pool if any accumulation (or depletion) is exactly balanced by another metabolite’s pool depletion (or accumulation), respectively. Some metabolites are not a part of any CM-pools as both their accumulation and depletion can be compensated via exchange with the extracellular medium. Finally, the accumulation or depletion of a metabolite cannot be re-balanced by perturbing the pool of other metabolites or via exchange with the medium (see Supplementary Materials 2 for the detailed *CMP-check* results). We found that the generation of CM-pool specific constraints scales poorly and becomes computationally inefficient for genome-scale models. Instead, we chose to indirectly satisfy all such CM-pools by simply projecting all RHS vector samples back to stoichiometric feasibility by minimizing the sum of violations (see *PULP* formulation in Section 2.3) while ensuring moiety and elemental balances.

Figure 6 provides a pictorial representation of the *PULP* optimization formulation. Randomly sampled RHS terms are projected onto feasible assignments for the RHS vector as dictated by network stoichiometry and reaction directionalities. This was implemented by adding slack variables to the randomly sampled RHS terms and employing as the objective function the minimization of the MW weighted sum of these slack variables. This weighting converts the projection to a mass-basis (i.e., gram gDW^−1^ h^−1^) from a molar-basis (i.e., mmol gDW^−1^ h^−1^), thereby preventing the RHS vector adjustment to bias the accumulation or depletion of metabolites with higher MWs. The same mass-basis was used to generate RHS random samples (see Section 2.3 for details on the sampling procedure).

**Figure 6.**
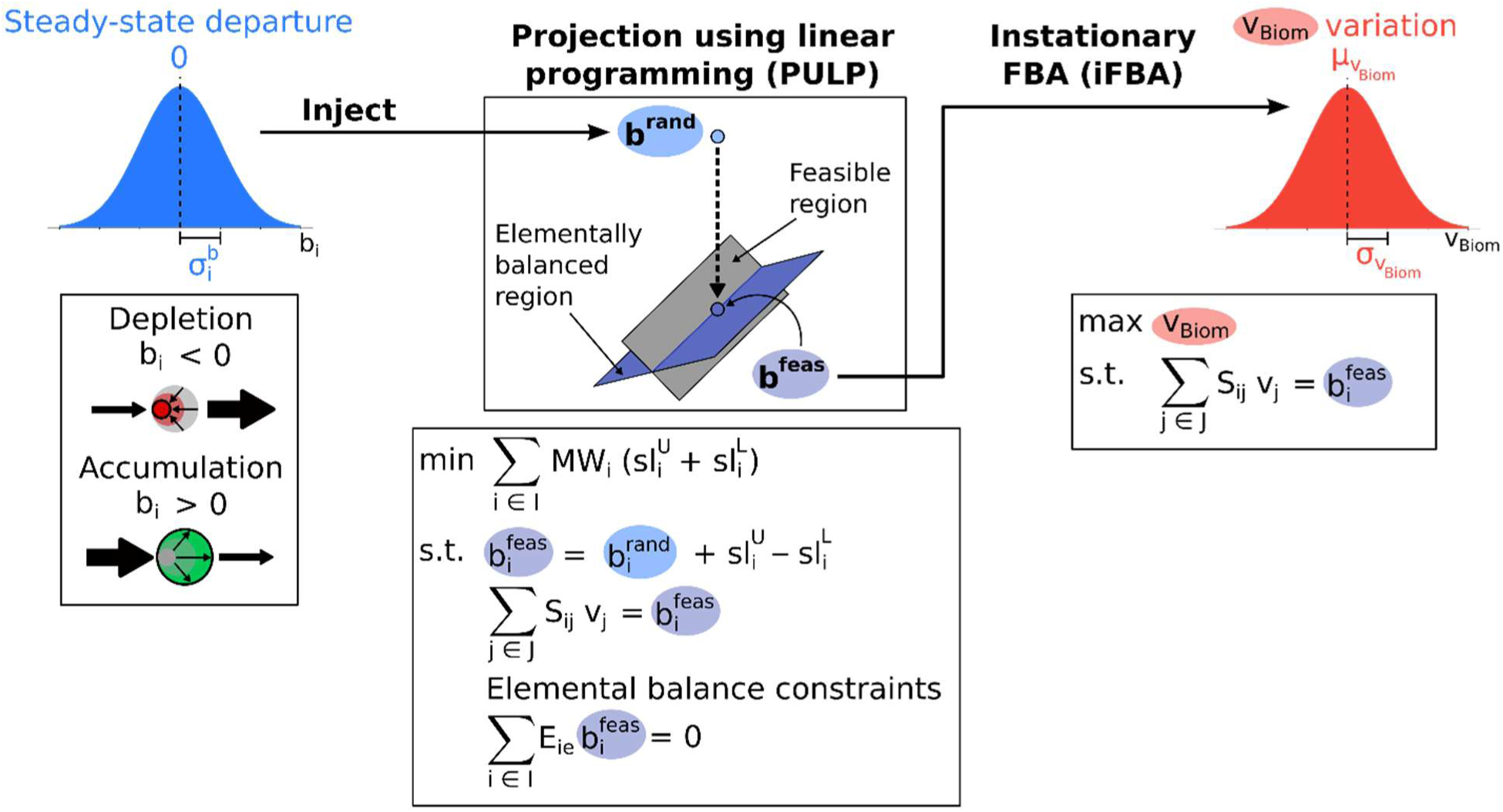
Overview of the method to sample and propagate uncertainty from metabolites unsteady-state. First, an ensemble of ***b***^***rand***^ vectors is generated by sampling from a normal distribution *𝒩*(0, *σ*^*b*^). Next, the optimization formulation *PULP* is used to project (one-at-a-time) the vector ***b***^***rand***^ onto the closest RHS vector ***b***^***feas***^ that satisfies both the conserved metabolite pools and elemental balances. These vectors ***b***^***feas***^ are used as inputs for *iFBA* to generate biomass yield predictions under RHS uncertainty.

Feasible RHS vectors ***b***^***feas***^ obtained from *PULP* are then used as inputs to the *iFBA* optimization (see Section 2.3) to calculate flux distributions and biomass yields and perform statistical analysis. Upon plotting the probability distributions of 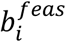, we find that they remain approximately normal for most metabolites (see Supplementary Text for a few example histograms). An important observation is that the ***b***^***feas***^ from *PULP* without the direct inclusion of the elemental balance constraints (see Section 2.3) introduces elemental imbalances to the *iFBA* output. This is because under steady-state FBA, all reactions carry out metabolite conversions that are automatically elementally balanced, assuming that the model stoichiometry is elementally balanced (Chan et al., 2017). However, in *iFBA* the departure vectors ***b***^***feas***^ introduce metabolite surpluses and/or deficits in a manner unconstrained by the model stoichiometric coefficients. The indirect incorporation of CM-pool constraints based on the *PULP* projection formulation ensures that the conservation of moieties (e.g., ACP, CoA, etc.) is maintained but elemental balances are not necessarily satisfied. This means that elemental balances must be directly imposed in the *PULP* formulation (see Section 2.3). To demonstrate what happens when the elemental balance constraints are not satisfied, we performed calculations using a modified version of *PULP* where we maximized growth under unsteady-state, both with and without the elemental balance constraints (see Figure 7A for an illustration). The upper bound of metabolite accumulation and depletion rates is set to 0.1% of glucose uptake rate, by mass. As expected, the cell can deplete some metabolite pools (i.e., 446) to (temporarily) grow faster by 5.4% while accumulating other metabolites (i.e., 431) (Figure 7B). This is achieved by inducing local imbalances around ATP and NAD(P)H producing reactions. For example, by depleting fructose-1,6P (fdp), glyceraldehyde-3P (g3p) and dihydroxyacetone phosphate (dhap) and simultaneously accumulating downstream metabolites, the cell can temporarily drive more flux through the payoff phase of glycolysis and yield up to 0.7% more ATP and 1.7% more NADH (see Figure 7C). However, lack of inclusion of elemental balances leads to a systematic depletion of metabolite pools (i.e., 1,085 depleted vs. 135 in surplus, respectively). Pool depletion is used as an alternative substrate source thus bypassing ATP dependent exchange reactions and reaching an erroneously predicted increase in biomass yield of 112.5% (Figure 7B). Therefore, care must be taken when interpreting unsteady-state FBA results because when casting the RHS vector as an unconstrained optimization variable, violations of the conserved metabolite pools and elemental balances can occur, leading to unrealizable flux distributions and an overprediction of the maximum possible biomass yield.

**Figure 7.**
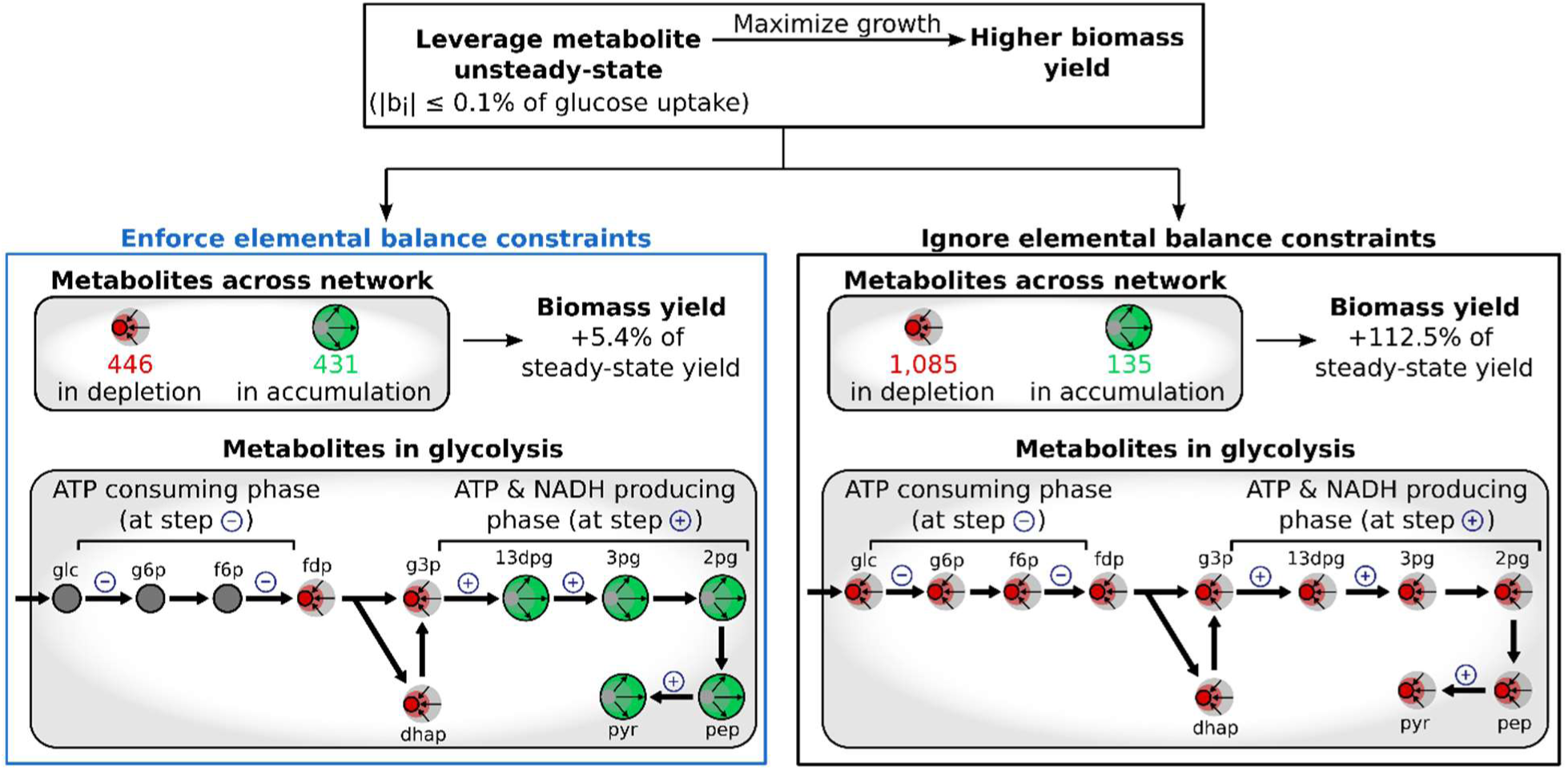
Maximization of biomass yield under metabolite unsteady-state for the two cases that enforce (left) or ignore (right) elemental balance constraints. Shown for each case are the numbers of metabolites depleting or accumulating, biomass yield change, and a closer look at the glycolysis pathway.

Neglecting the explicit introduction of elemental balance constraints during sampling using *PULP* introduces systematic elemental surplus which artificially boosts biomass yield. Figure 8 illustrates biomass yield predictions (i.e., cloud of blue dots) as a function of the percent elemental imbalance (C, N, P, and S) caused by the sampling without considering elemental balance constraints. Lack of consideration of carbon elemental balance causes most instances to be in overall carbon deficit as *iFBA* maximally drains carbon substrates towards biomass. Despite this negative balance, only a fraction of *iFBA* simulations lead to results that exceed steady-state FBA biomass yield. This is because metabolite imbalances cause biomass constituents to not be produced in the required ratios. The red vertical streak of points depicts the simulation when the elemental balance constraints are imposed. As expected, they always satisfy carbon balance and in the great majority of cases have a significantly decreased biomass yield, alluding to the largely detrimental effect of departing (even temporarily) from steady-state. Even though N and P are not limiting, as ammonium and phosphate uptake rates were left unconstrained, some enhancement of biomass yield as a consequence of their elemental imbalances were still observed (Figure 8). This is because N and P supplied from the growth medium require energy (i.e., NADPH or ATP) to be assimilated, which can be bypassed by violating elemental balance constraints and leveraging imbalances in intracellular metabolites. Note that for nitrogen incorporation through the carrier L-glutamate an NADPH molecule is required whereas for phosphate incorporation to a molecule an ATP molecule is typically required. Notably even though sulfur has a similar assimilation energy requirement via sulphate adenyltransferase requiring 2 ATPs and a GTP, no significant correlation between biomass yield and sulfur surplus was observed (see Figure 8). This is likely because sulfur accounts for only 0.8% of the biomass by weight compared to 49.3% for carbon, 15.1% for nitrogen and 9.2% for phosphorous.

**Figure 8.**
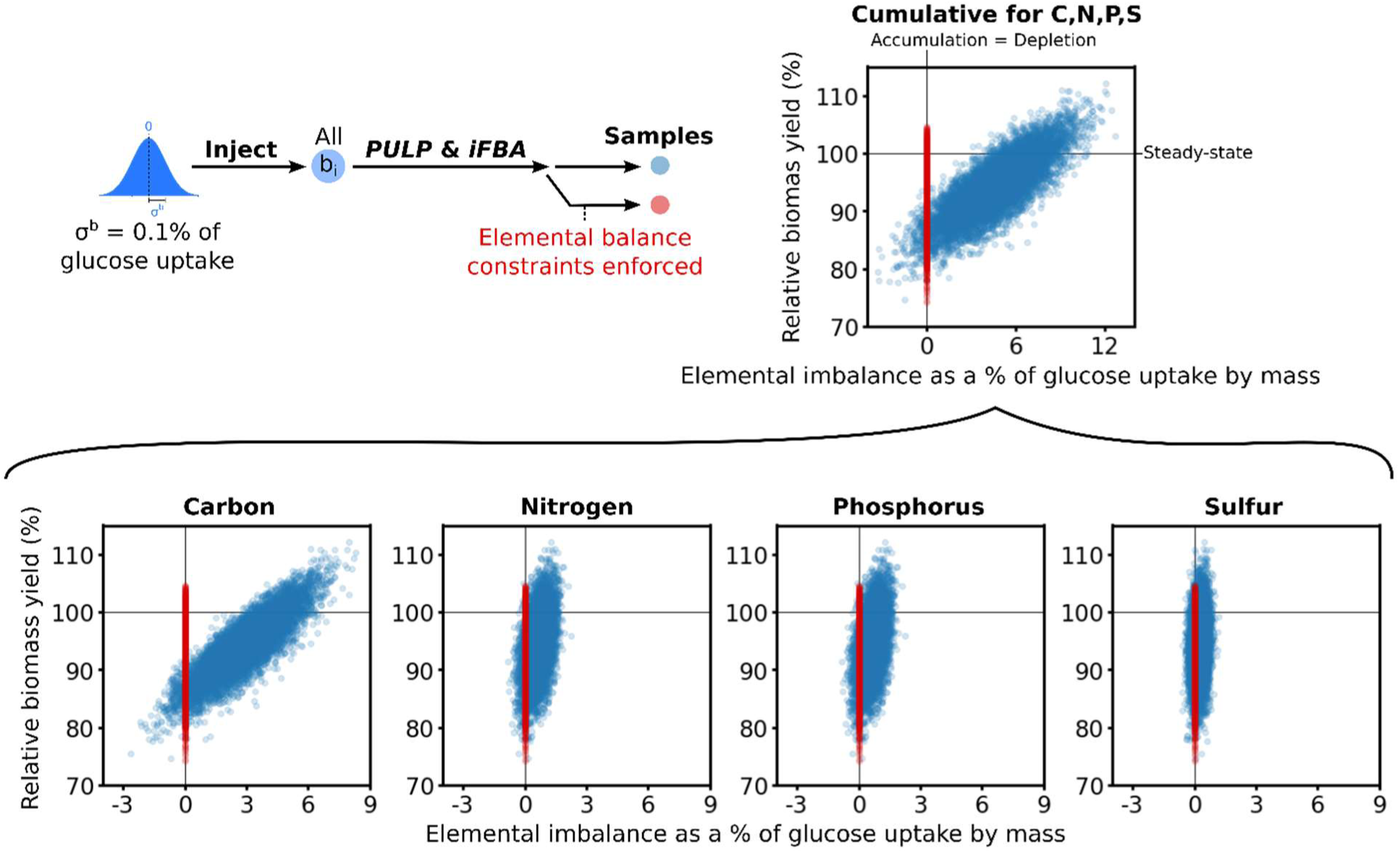
*iFBA* calculated biomass yields vs. incurred elemental imbalance as a % of the glucose uptake by mass upon ignoring (samples in blue) and enforcing (samples in red) elemental balance constraints.

Interestingly, we found that average biomass yield is much more sensitive to noise injection to the RHS vectors of the steady-state constraints than to the biomass composition coefficients. Even for an injected random noise with a standard deviation of just 0.1% of the glucose uptake rate (by mass) to the RHS, the average biomass yield for the population decreases by 7.1% from the steady-state yield (Figure 9). More substantial departures from steady-state attained by injecting noise that is 1% or 10% of the glucose uptake rate, leads to dramatic reductions of the average biomass yield by 67.9% or 94.8%, respectively. While it is possible for the cell to achieve faster growth by temporarily leveraging metabolite imbalances (see Figure 7 and 9), the effect is rather small and temporary since the depleting metabolite pools will likely be exhausted relatively quickly. Because there is a significant penalty in the biomass yield whenever even temporary excursions from steady-state are carried out in a random manner, a strong selection pressure is implied for cells to maintain exponential growth under steady-state. This further bolsters the steady-state assumption in FBA in predicting cellular phenotypes under growth pressure over a time span.

**Figure 9.**
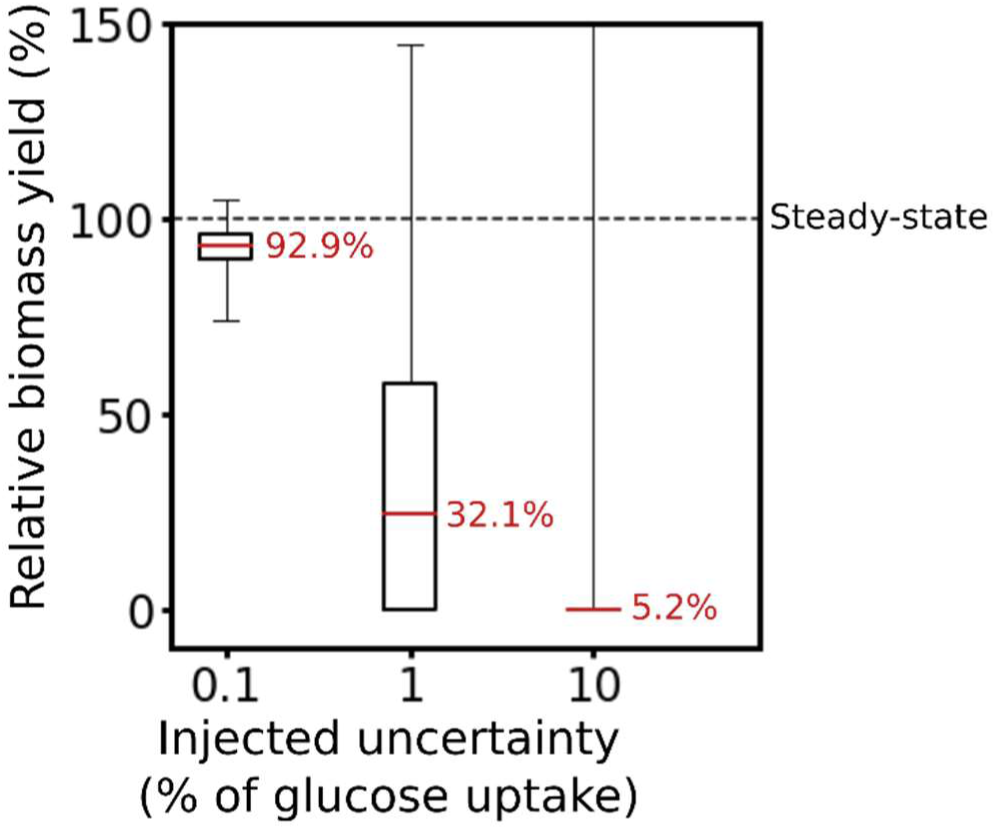
*iFBA*-calculated relative biomass yield when subjected to 0.1%, 1%, and 10% uncertainty in the RHS vector (as a mass % of glucose uptake rate). Biomass yield distributions are visualized with box plots and the numbers in red are the distribution means.

## 4. Discussion

In this work, we systematically quantified the effect of injecting random noise to biomass coefficient values and to the RHS of the metabolite balance equalities on FBA-predicted individual metabolic fluxes and biomass yield. Consistent with the robustness of biomass yield predictions by FBA, we found that random biomass coefficient uncertainty propagates weakly to the predicted biomass yield. This effect is overestimated by about 3-fold if the biomass coefficients are not property rescaled so that they correctly map to the MW of the biomass metabolite being 1 g mmol^−1^. In contrast, the effect on predicted individual metabolic fluxes can be much more pronounced, and in some cases amplified in comparison to the magnitude of the injected noise. Noise injected in the energy requirement component of the biomass reaction GAM has a proportionate effect on both FBA predicted biomass yield and metabolic fluxes, alluding to the importance of deriving an accurate estimate for GAM in metabolic model reconstruction and avoiding whenever possible direct porting from related organisms. Injection of noise on the RHS of balance constraints revealed the difficulty of recovering feasible flux distributions as metabolite pool and elemental balance constraints maintain strict requirements even under unsteady-state conditions. Metabolite unsteady-state causes dramatic reductions in the predicted average biomass yield even for very small temporary random departures from steady-state, pointing at a strong selection pressure to maintain steady-state during exponential growth. Conceptually, this implies that it is best for all biomass constituents to be available on a “just-in-time” basis to avoid growth arrests because a single required (core) biomass constituent is missing. Any potential benefit of temporarily boosting growth by creating a metabolite imbalance through depletion seems to cause a disproportionate reduction in the long run. The results obtained for both types of uncertainty argue in favor of the core assumptions of constant biomass composition and steady-state underpinning FBA.

This study revealed a number of best practices (and things to avoid) whenever assessing the impact of parametric uncertainty on FBA results. Any modification in the biomass reaction should be accompanied by checking for material balances in the lumped reactions (i.e., ATP hydrolysis for GAM), processes (i.e., DNA, RNA, and protein synthesis), and re-normalizing biomass MW to 1 g mmol^−1^. Similarly, updating biomass reaction formulations to account for new processes such as tRNA-charging of amino acids (Xavier et al., 2017) must be carried out so as mass and charge balances are conserved. An important observation from this study is that injected noise in biomass coefficients does not propagate equally to the biomass yield output, but it is dependent on the biosynthetic, energy, and cofactor costs for the corresponding biomass constituent. This establishes a prioritization sequence for experimental determination of various components of the biomass reaction. For example, amino acid composition and protein fraction should be prioritized in order to obtain accurate NADPH yield prediction whereas lipid species composition and fraction are important for accurate biomass yield prediction. Priority should also be given to macromolecular composition as their uncertainty propagation affects FBA-predictions to a larger extent than individual biomass coefficients. To this end, a pipeline was recently put forth (Simensen et al., 2021) to robustly measure macromolecular composition in *E. coli*.

Special considerations are also required when exploring departures from metabolite steady-states, as violating CM-pool equality constraints will cause the optimization problems to become infeasible. This can be a problem when incorporating experimental measurements of concentration changes to *iFBA* to quantify the RHS imbalance for metabolites that are parts of CM-pools. Missing or imprecise measurements could render the experimentally derived RHS vector unusable, and may require steady-state departures of unmeasured metabolite(s) to achieve a feasible solution (Bordbar et al., 2017). To this end, *PULP* can be a useful tool for projecting experimental measurements back to the feasible RHS solution space. Explicit inclusion of elemental balances within *PULP* is necessary to avoid elemental imbalances in the predicted flux distribution as under unsteady-state conditions the model stoichiometry cannot solely maintain elemental balancing leading to systematic depletions.

The methods developed here enable the prediction of flux phenotypes subject to an input of parametric uncertainty. They are also applicable to the study of systems that exhibit periodic variations in their metabolism (Andreas Angermayr et al., 2016) or the study of non-growing cells that only exhibit metabolite concentration fluctuations (Yurkovich et al., 2017). The assumption of normality for the injected uncertainty can be easily removed by generating samples conforming to any probability distribution. For example, one can assess what input probability distribution recovers the observed bimodal distributions of phenotypes (Elsafadi et al., 2016; Schmitz et al., 2017). Overall, the methods presented in this work could be used to further studies investigating metabolic flux and yield variation under parametric uncertainty in FBA.

## CReDIT authorship contribution statement

**Hoang V. Dinh**: Conceptualization, Methodology, Software, Validation, Formal analysis, Data Curation, Writing – Original Draft. **Debolina Sarkar**: Methodology, Software, Formal analysis, Writing – Review & Editing. **Costas D. Maranas**: Conceptualization, Methodology, Resources, Supervision, Writing – Review & Editing, Funding acquisition.

## Acknowledgment

We would like to thank Charles Foster (from The Pennsylvania State University) for a critical review of the manuscript. Computations for this research were performed on the Pennsylvania State University’s Institute for Computational and Data Sciences’ Roar supercomputer. This work was partially funded by the DOE Center for Advanced Bioenergy and Bioproducts Innovation (U.S. Department of Energy, Office of Science, Office of Biological and Environmental Research under Award Number DE-SC0018420). Any opinions, findings, and conclusions or recommendations expressed in this publication are those of the author(s) and do not necessarily reflect the views of the U.S. Department of Energy. Funding also provided by the DOE Office of Science, Office of Biological and Environmental Research (Award Number DE-SC0018260). Funding also provided by The Center for Bioenergy Innovation a U.S. Department of Energy Research Center supported by the Office of Biological and Environmental Research in the DOE Office of Science.

