## Supplementary Text for "Quantifying the propagation of parametric uncertainty on flux balance analysis"

### List of sections

S1. Standard deviation ratio (SDR) of flux sum

S2. Analysis of *CMP-check* results for in-pool metabolites

S3. Enumeration and analysis of minimal CM-pools

S3.1. CMP-find optimization formulation to enumerate minimal CM-pools

S3.2. Size and motif analysis for minimal CM-pools found with CMP-find

S3.3. Challenges in reconstruction of complete CM-pools from minimal CM-pools

S4. Example histograms for RHS terms projected by *PULP*

S5. Modified *PULP* formulation with growth maximization objective

### S1. Standard deviation ratio (SDR) of flux sum

SDR of flux sum, as compared to SDR of individual reaction fluxes, is a better metric to quantify the uncertainty of the net flux going from a metabolite node to another, because flux can equivalently go through all alternative pathways, leading to inflated SDR of individual reactions. Do note that in the particular case of glycoaldehyde degradation pathways, the flux sum metric is applicable even though the end metabolite nodes are different (i.e., malate or erythrose 4-phosphate). This is because the net glycoaldehyde degradation flux is extremely small (i.e., 0.006% of glucose uptake flux on average) thus the difference in the end nodes do not affect the flux vector as a whole. As expected, we found that SDR of the net flux (i.e., sum of fluxes) through all pathways was much lower compared to SDR of individual reactions/pathways (see Supplementary Materials 1 for the SDR of the “flux sums”).

**Table S1**. Alternative metabolic pathways activated under biomass composition and ATP maintenance uncertainty

| **Alternative modality** | **Reaction/Pathway** | **Reaction IDs*** |
| --- | --- | --- |
|  | ATP or GTP | |
| Cofactor | Adenylate kinase | ADK1 or ADK3 |
|  | ADP or GDP or UDP or CDP or dTDP |  |
|  | Pyruvate kinase | PYK or PYK2 or PYK3 or PYK4 or PYK6 |
|  | ATP or phosphoenolpyruvate |  |
|  | GDP to GTP | NDPK1 or PYK3 |
|  | UDP to UTP | NDPK2 or PYK2 |
|  | CDP to CTP | NDPK3 or PYK4 |
|  | dTDP to dTTP | NDPK4 or PYK6 |
|  | NADH or NADPH |  |
|  | FAD reductase | FADRx or FADRx2 |
|  | Quinone or menaquinone |  |
|  | Quinone monooxygenase | QMO2 or QMO3 |
|  | NADH dehydrogenase | NADH16pp or NADH17pp |
|  | Glycolate oxidase | GLYCTO2 or GLYCTO3 |
|  | L-Aspartate oxidase | ASPO3 or ASPO6 |
|  | Formate dehydrogenase | FDH4pp or FDH5pp |
|  | Formate + ATP or 10-formyl-THF |  |
|  | Glycineamide ribonucleotide (GAR) to 5’-phosphoribosylformylglycinamidine (FGAM) | GART or GARFT |
| Alternative transport shuttle moiety | Glycolate or glutamate or proline shuttling for Na^+^ uptake | (GLYCLTt2rpp and GLYCLTt4pp) or (GLUt2rpp and GLUt4pp) or (PROt2rpp and PROt4pp) |
| Alternative glucose uptake pathway | Glucose uptake via proton symport plus hexokinase or phosphotransferase system | (GLCt2pp and HEX1) or GLCptspp |
| Glycoaldehyde degradation | Glycoaldehyde to malate or erythrose 4-phosphate | (GCALDD and (GLYCTO2 or GLYCTO3) and MALS) or (4HTHRA and 4HTHRK and OHPBAT and PERD and E4PD) |

**^*^** “and” indicates reaction step relationship in a pathway, and “or” indicate alternative pathway relationship. Details about the reactions can be found in the *i*ML1515 model [1].

### S2. Analysis of *CMP-check* results for in-pool metabolites

We hereby provide an analysis on the metabolic pathway of the in-pool metabolites found via *CMP-check* (see main text, Section 2.3). The *CMP-check* outcomes for 1,536 intracellular metabolites in *E. coli* *i*ML1515 model are shown in Figure S1B, highlighting the in-pool and not-in-a-pool metabolites. When classifying by pathways, we identified that most of in-pool metabolites belong to select pathways (Figure S1C) (see Supplementary Materials 2 for pathway classification). For example, some cofactors and cell wall metabolites form a pool due to the lack of catabolic pathways and transport and exchange reactions. In inorganic ion transport and metabolism, some ions form a pool for only the depletion (i.e., $b_{i}$ < 0) direction due to a lack of export reactions. For other ion-associated metabolites, they contain conserved chemical moieties such as the aerobactin moiety participating as a chelating agent in iron sequestration. In alternate carbon metabolism, metabolites form a pool because they are disconnected from the metabolic network under glucose-utilizing conditions. In oxidative phosphorylation, a majority of metabolites are oxidized and reduced cofactors form a pool with each other.

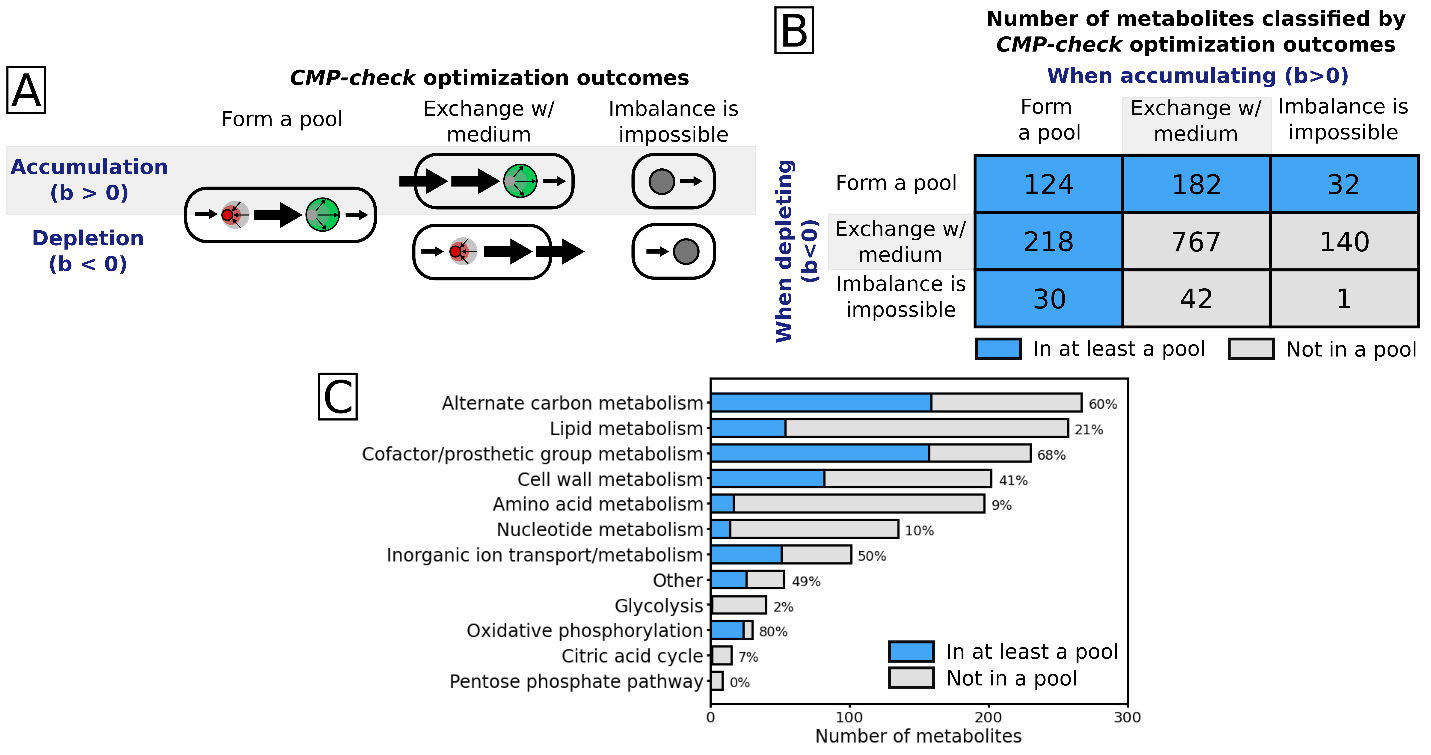

**Figure S1**. Pathway analysis of in-pool metabolites found via *CMP-check* optimization. (A) Visualization of all possible *CMP-check* optimization outcomes. (B) Statistics of *CMP-check* optimization outcomes with in-pool metabolites highlighted in blue and not-in-pool metabolites in grey. (C) Statistics of in-pool metabolites by metabolic pathway. Percentage of in-pool metabolites are shown.

### S3. Enumeration and analysis of minimal CM-pools

#### S3.1. CMP-find optimization formulation to enumerate minimal CM-pools

To find all in-pool metabolite pools we used integer cuts in the *CMP-check* formulation (see main text, section 2.4) with Boolean variables $y_{i}^{L}$ and $y_{i}^{U}$, which turn the respective slack variables $sl_{i}^{L}$ and $sl_{i}^{U}$ on (value of 1) or off (value of zero). The resulting formulation, called *CMP-find*, is as follows:

| $\min\sum_{i\in I,i\neq i^{*}} y_{i}^{U}+y_{i}^{L}$  $subject to$  $\sum_{j\in J} S_{i^{*}j}v_{j}=\epsilon$  $\sum_{j\in J} S_{ij}v_{j}={sl}_{i}^{U}-{sl}_{i}^{L}, \forall i\in I, i\neq i^{*}$  $0\leq sl_{i}^{U}\leq y_{i}^{U}M, \forall i\in I, i\neq i^{*}$  $0\leq sl_{i}^{L}\leq y_{i}^{L}M, \forall i\in I, i\neq i^{*}$  $y_{i}^{U}+ y_{i}^{L}\leq1, \forall i\in I, i\neq i^{*}$  $\sum_{i\in I_{k}^{P}} y_{i}^{L}+y_{i}^{U}<size\left( I_{k}^{P} \right), k=1,\ldots,K$  $v_{j}^{L}\leq v_{j}\leq v_{j}^{U}, \forall j\in J \backslash\{Biom\}$  $v_{Biom}=0$  $y_{i}^{L},y_{i}^{U}\in\left\{ 0,1 \right\}, \forall i\in I, i\neq i^{*}$ | (Eq. S1,  *CMP-find*) |
| --- | --- |

where $\sum_{i} y_{i}^{L}+y_{i}^{U}<size\left( I_{k}^{P} \right)$ is the integer cut constraint, and $I_{k}^{P}$ stores previously found minimal sets of metabolites that form a pool, indexed by $k=1,\ldots,K$. This representation excludes the previously found solutions and any other solutions whose sets of metabolites are supersets of previously found solutions. Mathematically, the integer cut excludes solutions in which in-pool metabolites’ $y_{i}^{L}$ or $y_{i}^{U}$ assume value of 1 without imposing any restrictions to the binary variables (i.e., $y_{i}^{L}$ and $y_{i}^{U}$) of other metabolites. In addition, combined with the integer cut, the objective function which minimizes the number of metabolites in a CM-pool allows *CMP-find* solutions corresponding to the minimal CM-pools to be found first. By definition, a minimal CM-pool is a minimal set of metabolites that have to be under unsteady-state simultaneously. By iteratively accumulating integer cuts, all metabolite(s) form a minimal pool with metabolite $i^{*}$ can be found until formulation *CMP-find* becomes infeasible, implying that all alternative minimal sets of metabolites that form a pool with $i^{*}$ are found. Descriptions for additional variables in *CMP-find* formulation can be found in the main text.

#### S3.2. Size and motif analysis for minimal CM-pools found with CMP-find

Using *CMP-find*, we found 48,846 minimal CM‑pools comprising of 2 to 6 metabolites (Figure S2A). All alternative solutions were found except for the depletion of the metabolite malcoame_c, for which 25,921 alternative solutions were found after the run time of a week. The total number of minimal CM-pools are large due to the explosion of possible depletion-accumulation combinations (visually described in Figure S2B). All of the pools describe a depletion-accumulation relationship, except for a case involving two metabolites (Figure S2D, size 2’s motifs 3) describing a co-accumulation relationship from nutrient in the media. Pools of larger sizes involve motifs where multiple conserved moieties are exchanged and in varied network topologies (Figure S2D). All minimal CM-pools found using *CMP-find* are provided in the Supplementary Materials 2.

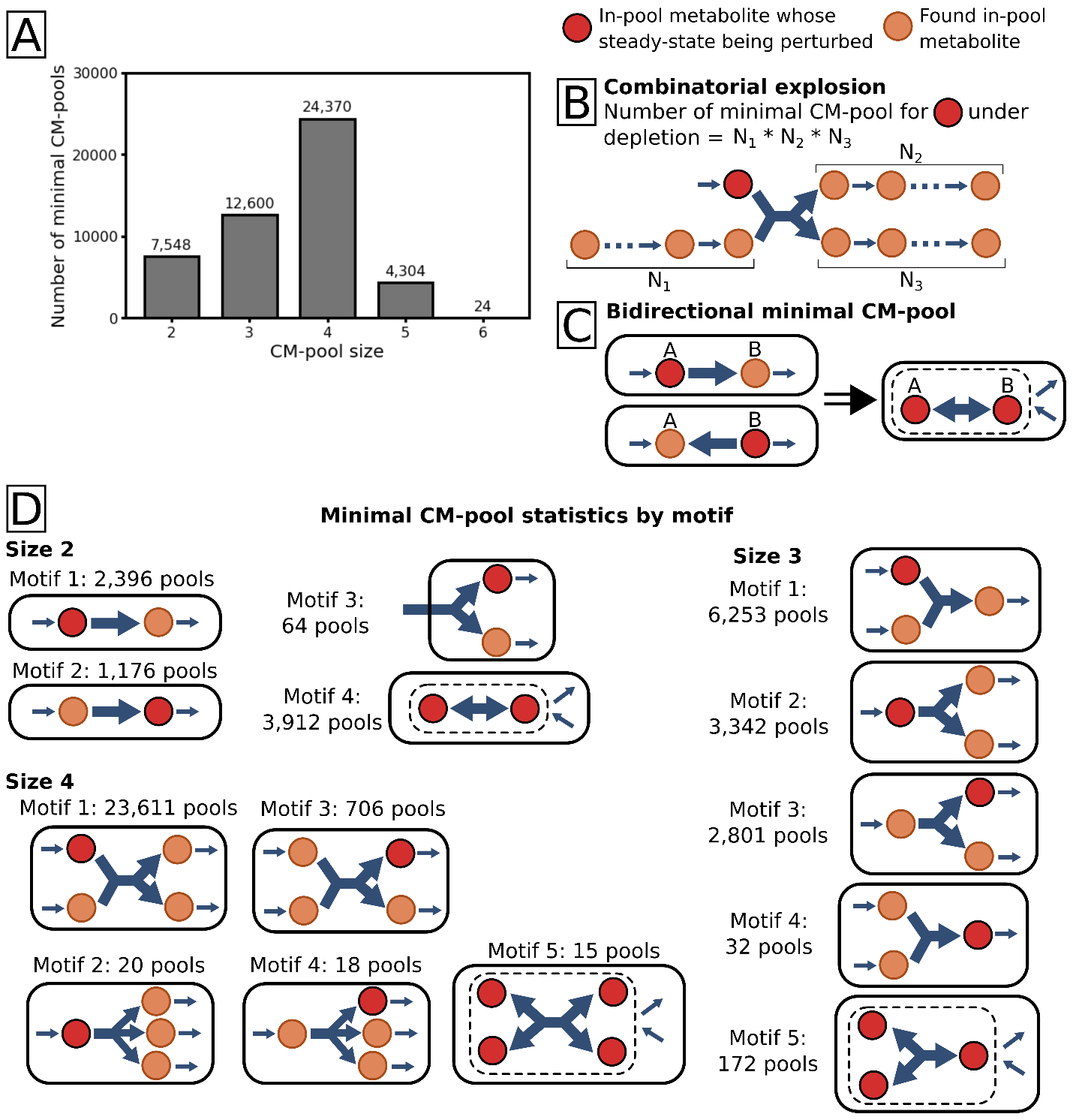

**Figure S2**. Minimal CM-pools statistics. (A) Number of minimal CM-pools found via *CMP-find* classified by pool size (i.e., number of metabolites per pool). (B) Visualization of combinatorial explosion leading to a large number of minimal CM-pools found. (C) Reconstruction of bidirectional pool from unidirectional pools. Depicted using a dashed border, metabolites in the bidirectional pool act as a single pool unit due to biochemical interconversion and influx and outflux correspond to the pool as a whole. (D) Visualization and number of minimal CM-pools for size 2-4. Size 5 and 6 pools follow similar motifs as present in size 4, but with a higher number of participating metabolites.

#### S3.3. Challenges in reconstruction of complete CM-pools from minimal CM-pools

The *CMP-check* outcomes for 1,536 intracellular metabolites in *E. coli* *i*ML1515 model are shown in Figure 5B (see Supplementary Materials 2 for the detailed *CMP-check* results). A key takeaway from *CMP-check* is that there are 1,291 metabolites (out of 1,536) whose RHSs can deviate from zero to either positive or negative direction. In other words, those metabolites are in-pool or not-in-pool for both accumulation and depletion. This means that there are a total of 2^1,291^ combinations of RHS signs. Only a subset of RHS sign combinations corresponds to feasible RHS vectors because CM-pools’ metabolites cannot be simultaneously under accumulation or depletion. We then extended *CMP-check* formulation to *CMP-find* by adding integer cuts to obtain all alternative solutions of CM-pools and found (incompletely) 48,846 minimal CM-pools after a run time of a week. A minimal CM-pool is a minimal set of metabolites that have to be under unsteady-state simultaneously.

While CM-pools resolve the RHS signs for metabolites within them, they do not resolve RHS signs for metabolites across the network simultaneously. This is because CM-pools’ metabolite sets are intersecting that requires method to resolve sign selection for the involved RHSs. For example, biotin CM-pool has metabolites that are also in CoA CM-pool and ACP CM-pool. In addition, because CoA is a precursor for ACP (see reaction ACPS1), CoA CM-pool and ACP CM-pool also intersect. Intersecting metabolites cannot be enumerated in a reasonable time period because 48,846 minimal CM-pools need to be aggregated into larger and complete CM-pools. In principle, aggregation of minimal CM-pools can be done by considering all metabolites in different minimal pools whose accumulation is coupled to the same particular metabolite’s depletion, or vice versa. The aggregation task is complicated by the fact that there are minimal CM-pools containing more than two metabolites and there are minimal CM-pools forming equality constraints with coefficients different than one. For example, in the isoprenyl pathway, there are the following minimal CM-pools for the depletion of isopentenyl diphosphate (as A) coupled with the accumulation of undecaprenyl diphosphate (as B) or octaprenyl diphosphate (as C): (i) $b_{A}+11b_{B}=0$, and (ii) $b_{A}+8b_{C}=0$. The coefficients correspond to the number of (conserved) isopentenyl groups in metabolites. In the Fe-S clusters biosynthesis pathway, the depletion of IscU-[4Fe4S] can be coupled to the accumulation of IscU protein and 4Fe4S metabolites simultaneously, based on the conservation of the two moieties (i.e., IscU and Fe-S cluster). Besides aggregating minimal CM-pools, another challenge in determining RHS sign combination is that some CM-pools can only associate with metabolite accumulation but not depletion (or vice versa) (see Figure 5B). In addition, if a CM-pool is active, the process of determining the RHS signs of metabolites in that pool can be cyclic. For example, let us consider a CM-pool where metabolite A’s depletion is coupled to metabolite B’s or C’s accumulation. If A is not being depleted, B or C do not need to be accumulated. If A is being depleted, then either B or C can still be depleted, but not together. Thus, the RHS sign selection process can be cyclic for A, B, and C as locking the RHS sign for one metabolite affects the RHS signs of others. This cyclic process issue is worsened due to the aforementioned issue of intersecting metabolites between CM-pools. Overall, through *CMP-check* and *CMP-find*, we found that finding CM-pools is computationally inefficient and the nature of metabolite pool and concentration coupling are complex at the genome scale.

### S4. Example histograms for RHS terms projected by *PULP*

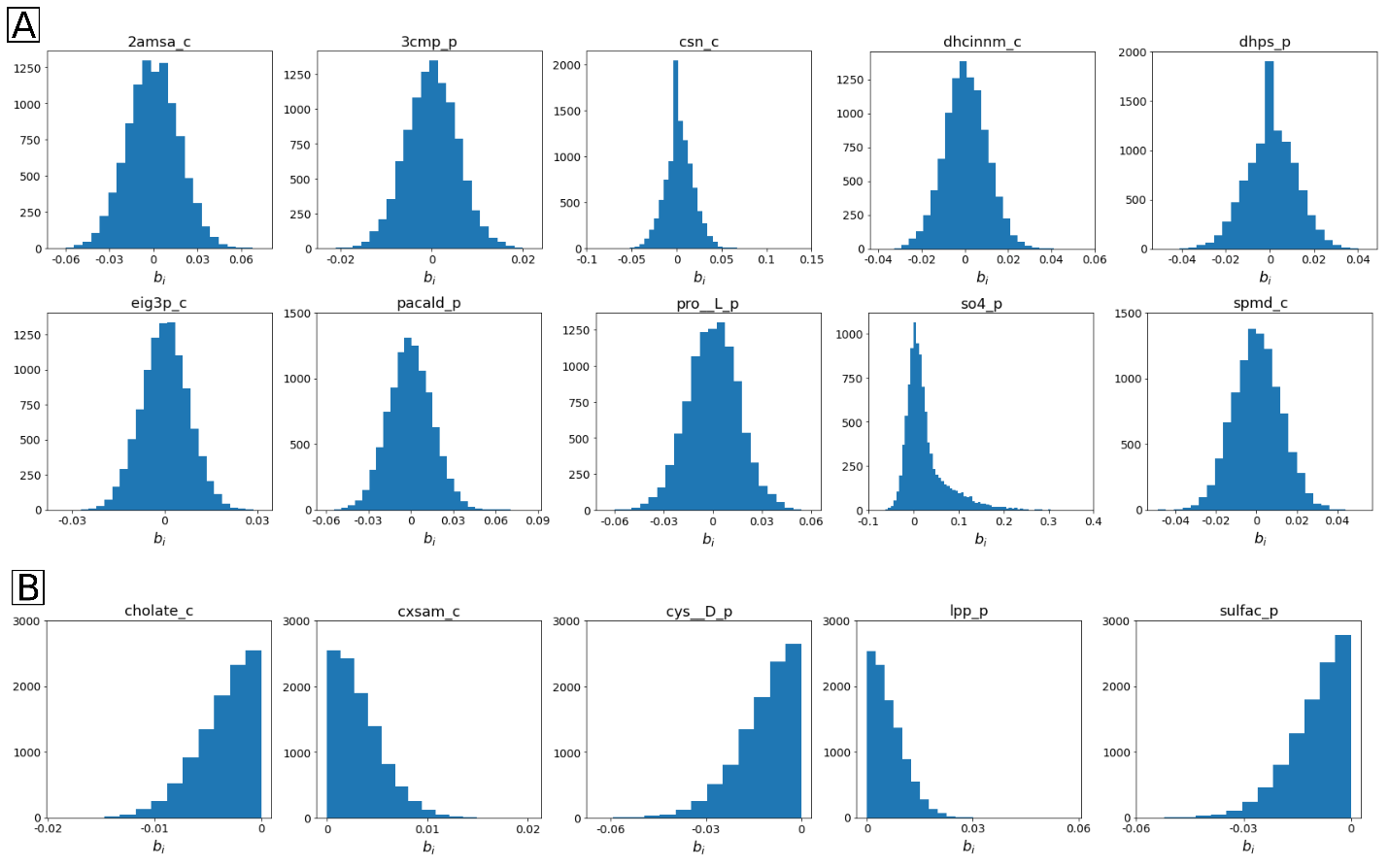

**Figure S3**. Example histograms for RHS term initially drawn from normal (A) and half-normal (B) distribution and adjusted by *PULP*. Ten (among 1,291) and five (among 245) metabolites were randomly selected among whose RHS terms drawn from normal and half-normal distribution, respectively. Metabolite ID is shown on top of each histogram.

### S5. Modified *PULP* formulation with growth maximization objective

To find the maximal growth yield subject to metabolite unsteady-state, we modify the *PULP* formulation as shown below:

| $\max v_{Biom}$  $subject to$  $\sum_{j\in J} S_{ij}v_{j}=b_{i}^{feas}, \forall i\in I$  $b_{i}^{feas}=sl_{i}^{U}-sl_{i}^{L}, \forall i\in I$  $0\leq sl_{i}^{U},sl_{i}^{L}\leq sl_{i}^{max}, \forall i\in I$  $v_{j}^{L}\leq v_{j}\leq v_{j}^{U}, \forall j\in J$  $\sum_{i\in I} E_{ie}b_{i}^{feas}=0, \forall e\mathcal{\in E}$ | (*PULP max growth*) |
| --- | --- |

where $sl_{i}^{max}$ is the upper bound (in mmol gDW^-1^ h^-1^) for the slack variables for metabolite $i$. All the upper bounds $sl_{i}^{max}$ is derived from the common mass basis in gram per gDW using Eq. 9 in the main text. Descriptions of additional symbols can be found in the main text.

To find the RHS vector subject to the maximal growth yield under metabolite unsteady-state, we perform a second round of optimization that minimize the MW-weighted sum of slack variables, as follows:

$$\min\sum_{i\in I} MW_{i} ({sl}_{i}^{U}+{sl}_{i}^{L})$$

$$subject to$$

$$\sum_{j\in J} S_{ij}v_{j}=b_{i}^{feas}, \forall i\in I$$

$$b_{i}^{feas}=sl_{i}^{U}-sl_{i}^{L}, \forall i\in I$$

$$0\leq sl_{i}^{U},sl_{i}^{L}\leq sl_{i}^{max}, \forall i\in I$$

$$v_{j}^{L}\leq v_{j}\leq v_{j}^{U}, \forall j\in J \backslash\{Biom\}$$

$$v_{Biom}=v_{Biom}^{max}$$

$$\sum_{i\in I} E_{ie}b_{i}^{feas}=0, \forall e\mathcal{\in E}$$

where $v_{Biom}^{max}$ is the maximal growth rate under metabolite unsteady-state found in the previous optimization of *PULP max growth*.

For the calculation without elemental balance constraints, the equations $\sum_{i\in I} E_{ie}b_{i}^{feas}=0, \forall e\mathcal{\in E}$, are removed from the optimization.
